# Excitation creates a distributed pattern of cortical suppression due to varied recurrent input

**DOI:** 10.1101/2022.08.31.505844

**Authors:** Jonathan F O’Rawe, Zhishang Zhou, Anna J Li, Paul K LaFosse, Hannah C Goldbach, Mark H Histed

**Author notes:** Department of Biological Structure, University of Washington, Seattle WA USA. National Eye Institute Intramural Program, National Institutes of Health, Bethesda MD USA.

## Abstract

Dense local, recurrent connections are a major feature of cortical circuits, yet how they affect neurons’ responses is unclear, with some studies reporting weak recurrent effects, some amplification, and others showing instead local suppression. Here, we show that optogenetic input to mouse V1 excitatory neurons generates salt-and-pepper patterns of both excitation and suppression. Responses in individual neurons are not strongly predicted by that neuron’s direct input. A balanced-state network model reconciles a set of diverse observations: the observed dynamics, suppressed responses, decoupling of input and output, and long tail of excited responses. The model shows recurrent excitatory-excitatory connections are strong and also variable across neurons. Together, these results demonstrate that excitatory recurrent connections can have major effects on cortical computations, by shaping and changing neurons’ responses to input.

## Introduction

The cerebral cortex of mammals is specialized into areas that perform different functions^1^. Animals from rodents to primates have several different visual cortical areas, each containing neurons with different types of selectivity^2–4^. In principle, these different representations in different visual areas could be created purely by feedforward mechanisms, where transformations happen via projections from one area or layer to the next, without outputs of a neuron feeding back (directly or indirectly) to influence that neuron’s activity. In fact, in a variety of artificial neural networks, much or all computation is provided by feedforward mechanisms^5^.

Yet in the brains of animals and humans, cortical recurrent connectivity is extensive. Most excitatory connections that a cortical neuron receives originate within a few hundred microns of their cell bodies^6–8^. Such recurrent connections can in principle have large effects on neural computation^9^, dramatically changing how cortical neurons respond to input.

How recurrent connections affect cortical computation is not fully understood, but important aspects of the structure of cortical recurrent connectivity have been determined. Some features of cortical network activity, such as irregular firing, are well-described by balanced-state models which assume strong recurrent coupling between excitatory and inhibitory neurons (either moderately strong, yielding ‘loose balance’, or very strong, yielding ‘tight balance’^10^). Work using inhibitory perturbations has shown that not just excitatory-inhibitory connectivity is strong, but the average excitatory-excitatory connectivity is strong as well. More precisely, cortical recurrent excitatory coupling is strong enough that the excitatory network is unstable and self-amplifying, a phenomenon described by inhibition-stabilized network models (ISNs)^11–14^.

While some consensus has developed on these average cortical connectivity properties (but see^15^), the effect recurrent connections have on transforming sensory or input signals has been less clear. For example, some recent studies have shown that certain patterns of excitatory input can be amplified by the cortical network^16,17^, consistent with some theoretical predictions^18,19^. On the other hand, however, some studies have shown that nearby neurons can be substantially suppressed by stimulation that excites a single or a small ensemble of excitatory cortical neurons^20,21^. How excitatory and inhibitory neurons might interact through recurrent connections to create such suppression has not been determined.

Here, to understand how cortical neurons’ responses are shaped by the cortical recurrent network, we stimulate excitatory cells in the visual cortex optogenetically and record responses of local neurons with electrophysiology and two-photon imaging. First, we find that stimulation of excitatory cells leads to a salt-and-pepper pattern of local suppression, consistent with the pattern of excited and suppressed cells produced when animals see a strong visual stimulus. To understand how this suppression effect might arise from cortical recurrent circuitry, we examine both the patterns of firing rate changes and the dynamics of responses. Recent theoretical work has shown that cortical visual responses can be “reshuffled” by additional excitatory input^22^ — that is, strong average recurrent coupling allows individual neurons’ firing to change significantly in response to input while the distribution of population activity is little-changed^23,24^. We implement this scenario in a conductance-based simulation and find that it can explain the suppression we observe. In addition, our data is consistent with substantial variability in local recurrent connectivity, with some neurons receiving large net recurrent excitation and others smaller or net suppressive recurrent input. Our results go beyond prior work that found strong *average* recurrent connectivity, showing that *variance* in excitatory-excitatory connectivity must also be substantial, and further show that this variance in recurrent connectivity can decouple neurons’ firing rate responses from the direct input they receive.

The suppression we observe during excitatory cell stimulation occurs in individual cells, but the mean response is elevated. This increase in mean, however, seems at odds with the prior finding that single-cell stimulation leads to inhibition on average^20^. To resolve this, we simulate the effect of single cell stimulation and find that the difference in the two results can be explained by the activation state of the cortical network. Increasing activity in the network with visual stimulation results in a slight decrease in mean responses to stimulation, showing the prior results and our current results can be described in the same model framework.

Thus, a balanced-state cortical model, with strong average coupling and variability in recurrent connectivity, explains many features of our data, including dynamics and neural response distributions. These results show how cortical neural suppression can be generated from excitatory input: variability in recurrent input means that firing rate responses are decoupled from (are only weakly affected by) the level of excitatory input we provide to that cell. This arises because much of the input a cell receives comes from recurrent sources. Because recurrent input varies from cell to cell, the result is many excited cells, but also a substantial number of suppressed neurons.

## Results

### Strong visual input leads to salt-and-pepper distributed suppression in primary visual cortex

We first measured local patterns of suppression in visual cortex in response to visual stimuli. We presented small high-contrast visual stimuli to headfixed mice while measuring activity in V1 layer 2/3 neurons via two-photon imaging (Fig. 1A). We expressed GCaMP7s in all neurons via viral injection (AAV-hSyn-GCaMP). Animals were kept awake and in an alert state^25^ with occasional drops of water reward.

**Figure 1:**
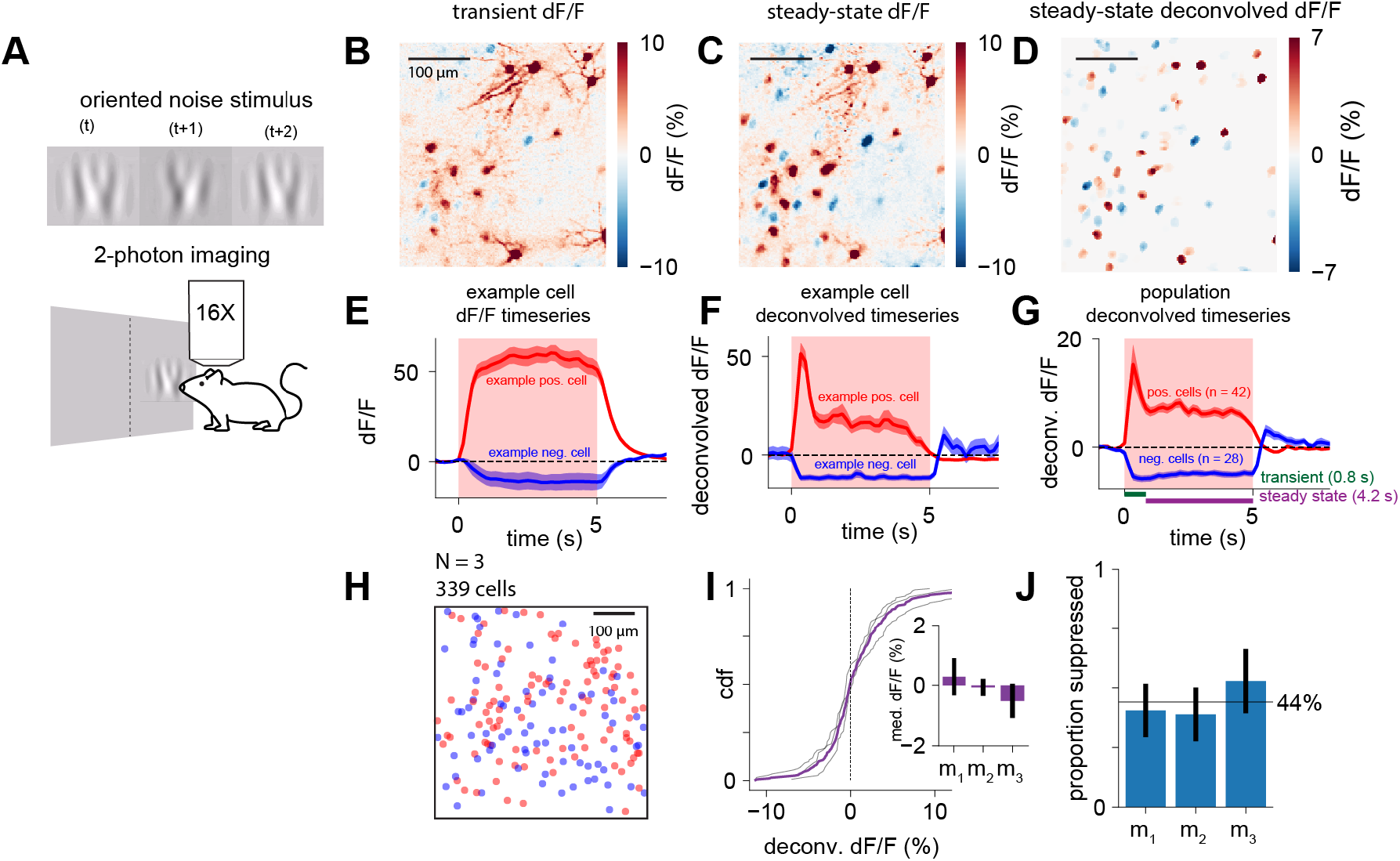
V1 neurons show salt-and-pepper suppression to strong visual stimuli. **(A)** Experimental setup. Awake mice viewed a small (15 degree diameter) visual stimulus with rapidly changing frames of oriented noise (Methods). **(B)** Example 2-photon imaging data from layer 2/3 of V1 in response to the stimulus, during the transient and **(C)** steady-state periods. Time intervals used for averaging in (B-D) displayed in green and purple in (G). Intermixed (salt-and-pepper) elevated and suppressed responses emerge during the steady-state period. **(D)** Deconvolved responses from (C), projected onto segmented cell masks (Methods). **(E)** Example dF/F trace for one elevated and one suppressed cell. Shaded regions: SEM across trials. Shaded red: optogenetic stimulation duration. **(F)** Deconvolution of the traces in (E) reveals an initial transient period and then a steady-state response. **(G)** Average response for all elevated and suppressed cells in (B-D, N = 1, pos. neurons = 42, neg. neurons = 28)**. (H**) Spatial distribution of elevated (red) and suppressed (blue) cells collapsed across animals (N = 3; 339 neurons), showing random distribution of neurons across the cortex (statistical analysis; Fig. S1F-H). **(I)** Visual response amplitudes are similar across animals. Thin lines: CDFs for individual animals, thick line: population CDF. Inset: medians are near zero, m_1_-m_3_: individual animals, error bars: ± SEM. **(J)** Proportion of cells suppressed in each mouse. Error bars: Wilson score 95% confidence intervals. Black line: group mean (44% ± 7%). See also Fig. S1.

We imaged responses to two types of high-contrast visual stimuli, a fast-changing stimulus designed to minimize adaptation (“oriented noise”, Fig. 1A)^26–28^ and a drifting grating (Fig. S1D). We found a salt-and-pepper mix of suppressed and excited cells (Fig. 1B,C), with suppression stronger after the initial stimulus response (Fig. 1C). In other words, in response to both types of visual stimuli, we found some cells that responded with strongly elevated steady-state responses, and other cells that showed suppressed responses (Fig. 1C–F, Fig. S1A-E).

Deconvolving fluorescence responses to yield a proxy measure of spike rate confirmed this salt-and-pepper pattern, with substantial numbers of suppressed and excited neurons intermingled (Fig. 1D). The deconvolution revealed an initial transient response in excited cells (Fig 1F,G), followed by either an elevated or suppressed steady state.

We confirmed that the spatial distribution of elevated and suppressed neurons was randomly scattered across the cortex (Fig. 1H). We found our data was consistent with random scatter (data vs 2d Poisson process model for spatial randomness, p > 0.05, Bonferroni correction, Fig. S1F-H).

The viral expression strategy we used for these experiments results in both excitatory and inhibitory neurons that express GCaMP. However, the large fraction of suppressed neurons (Fig. 1D,H-J; proportion suppressed 44% ± 7%, N=3 animals, mean ± standard error) implies that it is not that a group of inhibitory neurons was suppressed by stimulation, but that many excitatory neurons were suppressed. Below, we confirm with electrophysiology and imaging that optogenetic excitatory input produces suppression in many excitatory cells.

### Optogenetic excitatory drive also results in sparse and distributed suppression

To examine the influence of recurrent excitatory-inhibitory circuits on local response properties, we next measured V1 responses while optogenetically stimulating excitatory cortical cells (Fig. 2A). Direct stimulation allows us to exclude some feedforward mechanisms for suppression — for example, to argue against the possibility that cortical suppression is generated principally by suppression of thalamic inputs^29^.

**Figure 2:**
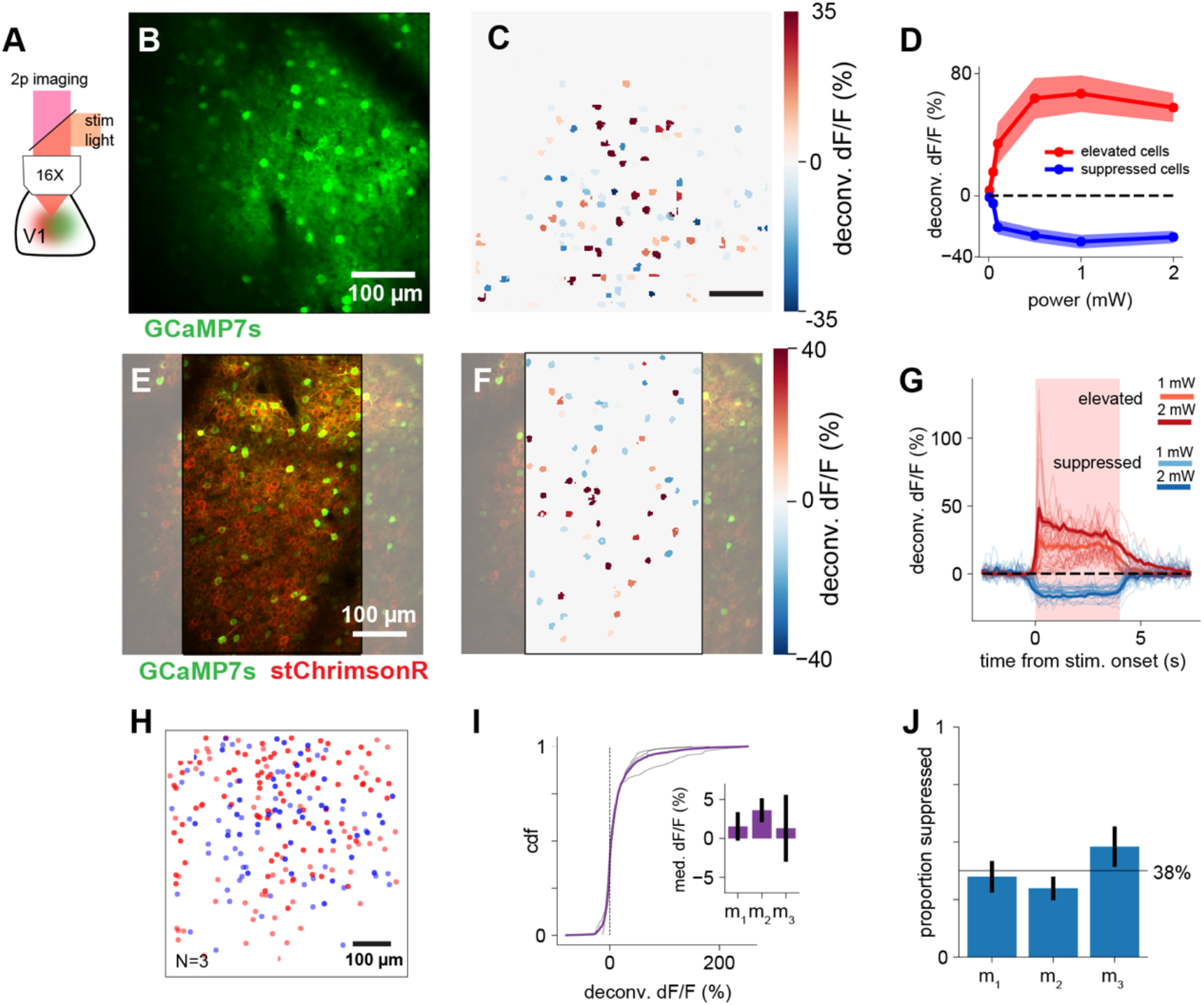
Salt-and-pepper elevation and suppression to optogenetic excitation. **(A)** Experimental setup, using two-photon imaging (GCaMP7s, all cells, 920nm) and optogenetic excitation of excitatory neurons (stChrimsonR, 595nm). **(B)** Example field of view. **(C)** Deconvolved steady-state response (scaled to match dF/F %) to optogenetic stimulation (200 ms duration) from (B). Red: elevation of firing rate relative to baseline, blue: suppression. **(D)** Increasing power leads to stronger elevation and suppression (steady-state response) in their respective populations. Shaded region: SEM across cells. **(E)** Field of view from an example animal stimulated with long (4 sec) optogenetic pulses; stimulation during imaging flyback (Methods). Gray: areas omitted from analysis to exclude stimulation artifact. **(F)** Deconvolved response to stimulation, conventions same as (C). **(G)** Population timecourses for cells in (F). Red region: optogenetic stimulation period. Steady-state response averaging period: 200-3750 ms. Light lines: individual cell traces, heavy lines: population averages. Shaded region (largely obscured by thick lines): SEM across cells. **(H)** Spatial distribution of elevated (red) and suppressed (blue) cells collapsed across all animals (N = 3), same conventions as Fig 1H. Statistical analysis: Fig. S1F-H. **(I**) Optogenetic response amplitudes are similar across animals. Conventions as in Fig. 1I**. (J)** Proportion of cells suppressed by optogenetic stimulation in each mouse. Error bars: Wilson score 95% confidence intervals. Black line: group mean (38% ± 8%).

We injected a Cre-dependent excitatory opsin (soma-targeted ChrimsonR, or stChrimsonR) in layer 2/3 of a mouse expressing Cre in excitatory neurons only (*Emx1-Cre*^30^), and expressed GCaMP7s in all neurons with a second virus (AAV-hSyn-GCaMP7s) (Fig. 2B,E).

With optogenetic stimulation we also found a clear salt-and-pepper distribution of elevated and suppressed responses (Fig. 2C,F,H; short stimulation pulses Fig. 2B-D, long pulses with imaging of steady-state during stimulation, Fig. 2E-G). Neural responses to stimulation increase as power increases (Fig. 2D; asymptote may be due to opsin saturation.) As in the case of visual responses, we confirmed that the spatial patterns of responses were compatible with random scattering (all p’s > 0.05, Fig. S1F-H). The proportion of suppressed neurons with optogenetic stimulation (Fig. 2I,J; 38% ± 8%, mean ± SEM) was comparable to that seen with visual stimulation (Fig. 1IJ). These optogenetic data suggest that the network is being driven to a new steady state or fixed point by input. While there was a slight decay in the excited population’s response at high power (perhaps due to network effects, spike rate adaptation, or opsin desensitization), at moderate stimulation power (1 mW, Fig. 2G), deconvolved firing rates are largely constant while stimulation is on.

We confirmed the opsin we used was expressed only in excitatory cells using fluorescence in-situ hybridization. We labeled excitatory, inhibitory, and stChrimsonR-expressing neurons (RNAScope, ACD Inc; Fig. S2A,B). Excitatory neurons expressed the opsin (Fig. 2I), but as expected for AAV expression^31^, not all excitatory neurons were opsin-positive (59%, N = 115/195, Wilson score 95% CI: [52.0%,65.7%], Fig. S2A). None of the inhibitory neurons (24% of neurons in the sample, N = 62/257) showed expression of the opsin (Fig. S2B).

The two-photon imaging experiments showed a salt-and-pepper pattern of excitation and suppression within the imaging fields of view. To examine whether this salt-and-pepper pattern exists at larger distances from the stimulation site, we used electrophysiology. We recorded neural responses to stChrimsonR stimulation using a silicon electrode array (Fig. 3A,E) and the same viral strategy for opsin expression as we used with imaging.

**Figure 3:**
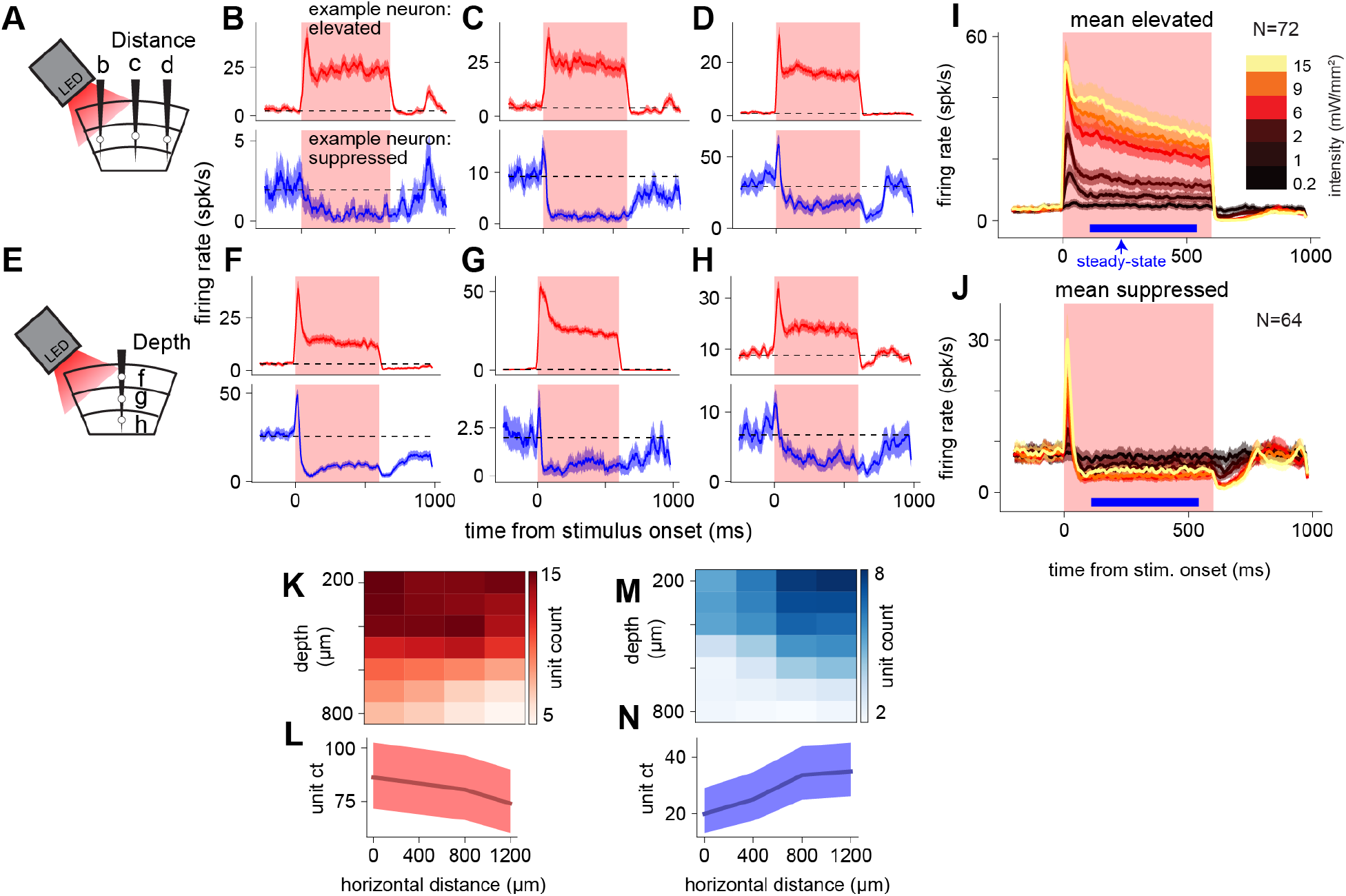
Stimulation of V1 excitatory neurons yields salt-and-pepper organization across the cortex. **(A)** Neural responses recorded across the cortex. Recordings in vivo from awake mice. **(B, C, D)** Example neurons at three distances from stimulation light (0 µm, 400 µm, 1200 µm), showing elevated and suppressed cells at all distances. **(E)** Neural responses recorded through cortical depth. **(F, G, H)** Example neurons recorded at three depths (250 µm, 550 µm, and 800 µm), showing elevated and suppressed cells at different depths. **(I)** Population average timecourses of elevated cells. Blue bar: interval for steady-state rate calculation. Shaded regions: SEM across cells. **(J)** Population time courses of suppressed cells, same conventions as (I). **(K)** Counts of elevated units (single and multi-units) by distance and depth, smoothed with a Gaussian kernel for display. **(L)** Distribution of elevated steady-state responses across horizontal distance, summed across depth. Shaded region: Wilson score 95% CIs. Note lower limit of y-axis not zero. **(M-N)** Same as (K-L), but for units with suppressed steady-state responses. See also Fig. S2.

We found both elevation and suppression across all distances (Fig. 3B–D) and depths (Fig. 3F– H) from the stimulation site, suggesting a similar salt-and-pepper organization of elevated and suppressed cells extends over distance. Across the population of recorded neurons, 56.6% (77 of 136) showed an elevated steady-state response to the optogenetic stimulation, and 36.0% (49 of 136) showed a suppressed steady-state response, comparable to our two-photon measurements (Fig. 2). Both elevated and suppressed cells on average showed an initial (positive) transient followed by a (positive or negative) steady-state response (Fig. 3I,J).

The electrophysiological recordings show similar dynamics as the deconvolved imaging timecourses (Fig. 2H), except for one feature: the recordings show an initial brief positive transient in the suppressed cells (Fig. 3B–D, F–H, blue lines; Fig. 3J) not just in the elevated cells as in the imaging data. This transient is likely concealed in the imaging data due to the slower timescale of imaging. The imaging frame rate (30 Hz; 33 ms frames) is slower than the transient, so within one frame the positive transient would be averaged with suppression, yielding a result near zero. In the case of elevated cells, the positive transient is averaged with an elevated steady state, and so the response in that frame remains positive.

### Global spatial patterns arise from trends in local salt-and-pepper suppression

The neurophysiology data showed some evidence of a larger-scale organization on top of the local salt-and-pepper distribution of elevation and suppression. Over distances of more than a millimeter from the stimulation site, we found that the number of elevated units gradually decreased (Fig. 3K,L; Pearson’s chi-squared test, χ^2^ = 51.31, df = 3, p < 0.001) and the number of suppressed units gradually increased (Fig. 3M,N; χ^2^ = 44.83, df = 3, p < 0.001; see Fig. S2E-H for unit counts as a proportion of total units). There was also a similar trend in neurons’ firing rates (Fig. S2I,J). Elevated single units showed less elevated firing rate with distance from the stimulation site, and suppressed single units showed more suppression with distance from the stimulation site, though the linear trend between distance and population response was stronger in unit counts than in average population firing rates (Pearson’s r = -0.11, df = 29, p = 0.56, Pearson’s r = 0.32, df = 46, p < 0.05; Fig. S2I,J). Notably, however, the number of elevated neurons did not go to zero even at 1.2 mm from the stimulation site: only the relative numbers of elevated and suppressed neurons changed. This suggests that the salt-and-pepper organization we saw with imaging persists across the cortex.

The trends over distance we saw with physiology, however, give only a partial view into how population responses varied with distance from the stimulation site. To measure the extent of suppression across the cortex, we turned to widefield, mesoscale calcium imaging. For these experiments, we expressed GCaMP in all excitatory cells using a genetic mouse line (to maximize consistency of GCaMP expression across cortical distance; Fig. 4A; Ai148::Cux2-CreERT2, or Ai162::Cux2-CreERT2, see Methods). We restricted expression of stChrimsonR to excitatory cells using the CamKIIa promoter (AAV-CamKIIa-stChrimsonR) and stimulated while simultaneously imaging responses.

**Figure 4:**
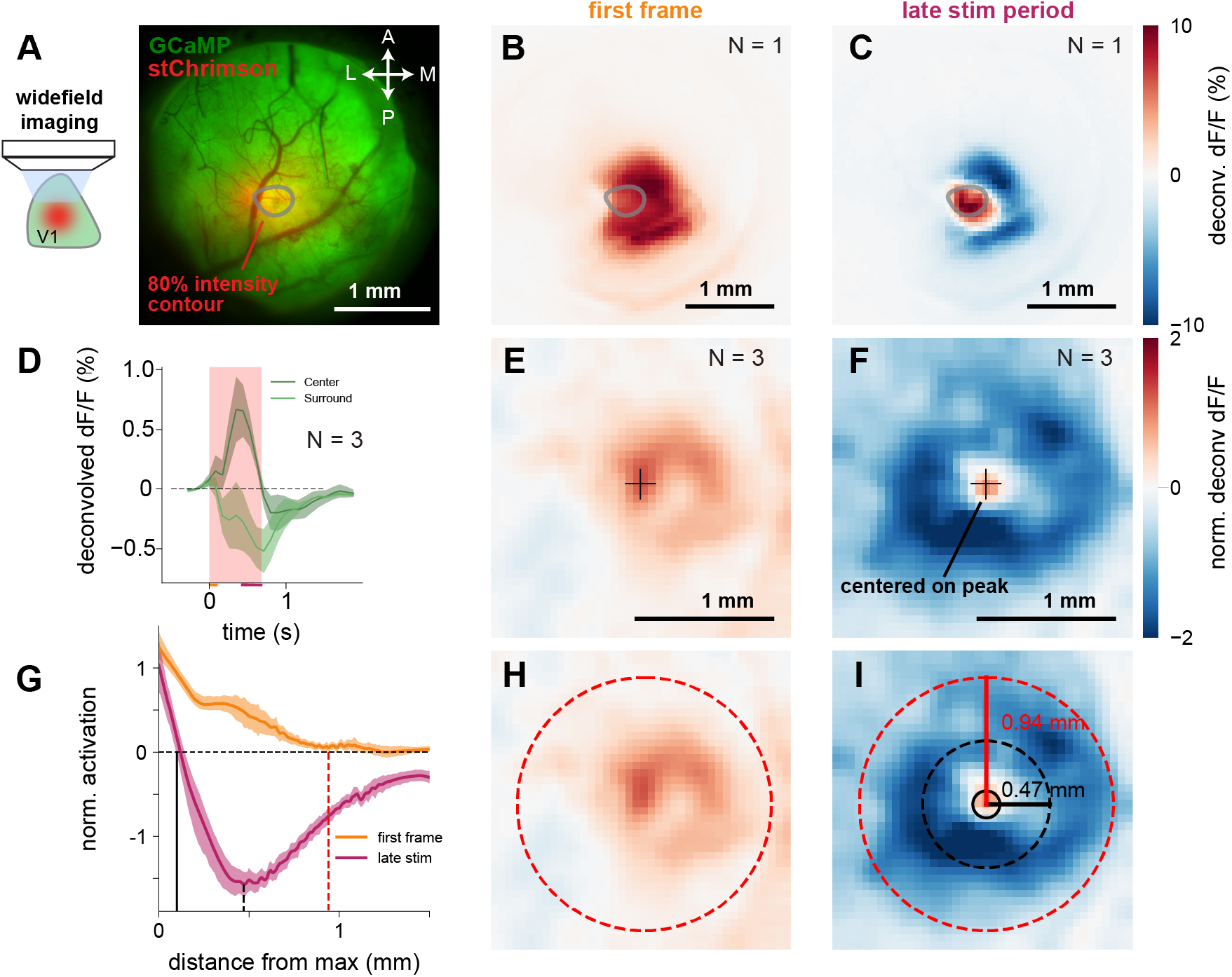
Widefield imaging of excitatory neurons shows average center-surround organization during steady-state periods. **(A)** Experimental setup: stChrimsonR in excitatory neurons via viral transfection (AAV-CamKIIa-stChrimsonR), expression of GCaMP via mouse line (either Ai148::Cux2-creERT2, GCaMP6f, or Ai162::Cux2-creERT2, GCaMP6s; induced with tamoxifen as adult; Methods). Right: imaging field of view for one animal. **(B-C)** Mean deconvolved response (see Fig. S3) during first frame (B) and during the late stimulation period (C) in an example animal (Fiber for light delivery slightly obstructs the imaging field, see Fig. S4D-F). **(D)** Average response to stimulation over time (N = 3). Red shaded region: stimulation period, orange bar: first frame, maroon bar: late stimulation (steady-state) time period. **(E,F)** Average responses, N = 3 animals. Responses for each animal were aligned spatially to the peak during the late stimulation period (Methods), smoothed for visualization. **(G)** Response as a function of distance, averaged from data in E, F. Smoothing: LOWESS. Shaded regions: bootstrapped 95% CIs. Vertical lines: zero crossings and inflection points. Zero crossings defined by shortest distance at which 95% CI included zero. Black lines: late stimulation period. Solid black: first zero crossing, dotted black: local minimum. Red dashed line: early response, first zero crossing. **(H, I)** Same as (E,F) but with superimposed circles whose radii correspond to lines in (G). See also Figs. S3-4.

We saw clear spatial patterns in widefield imaging, broadly consistent with the spatial trends we saw in the electrophysiology data. During the initial frame of stimulation (∼7 Hz imaging, 140 ms frame period), we saw an increase in activity both at the center of the stimulation light and extending some distance outside the center of expression (Fig. 4B,E,H).

A center-surround pattern emerged later in the stimulation pulse (Fig. 4C,F,I) consistent with the large-scale patterns in the electrophysiological recordings. The area with maximum stChrimsonR expression continued to show an elevated response, while a donut-shaped region around it was suppressed (see Fig. S4G-I for spatiotemporal response). The activated area in the center reflected the area of expression, measured with fluorescence imaging of the cortical surface (Fig. S4A-F). To examine these timecourses (Fig. 4D), we deconvolved imaging responses to yield approximations to spike rate changes. We compared several different deconvolution methods and found suppression in all cases (Fig. S3). The suppression was strongest about 500 µm from the center of our laser stimulus, and extended over 1 mm from the stimulation center (Fig. 4 G–I). In electrophysiology, the number of suppressed cells increases by a factor of two over approximately this distance (Fig. 3K-N), and therefore the increased number of suppressed individual neurons may be the substrate for the suppression in this imaging data.

In summary, the physiology and imaging data together support the idea that suppressed and elevated neurons are locally organized in a salt-and-pepper pattern, and that the proportion of suppressed to elevated neurons increases with distance from the stimulation site. This change in the proportion of suppressed cells results in a center-surround pattern that can be seen with population-level imaging, with net suppression in excitatory cells emerging, after an initial positive transient, about 500 µm away from the stimulation site.

### Response dynamics support a balanced-state excitatory-inhibitory network that is driven to a new steady state by input

If suppression is due to local recurrent network effects, we would expect excitatory cells to be recruited first by stimulation, and then inhibitory cells should receive inputs from excitatory cells and respond slightly later. After this first few milliseconds, balanced-state models predict that excitatory and inhibitory cells should later show similar response distributions^10,11,32,33^. This is in contrast to weakly-coupled models, or a feedforward inhibition framework, where excitatory and inhibitory populations change firing rates in opposite directions: that is, input drives inhibitory cells to increase their rates, inhibiting excitatory cells, which then decrease their rates.

Our data supports the balanced-state recurrent model (Fig. 5A) — we saw differences in excitatory and inhibitory responses in the first few milliseconds, but at later times distributions of excitatory and inhibitory rates were similar.

**Figure 5:**
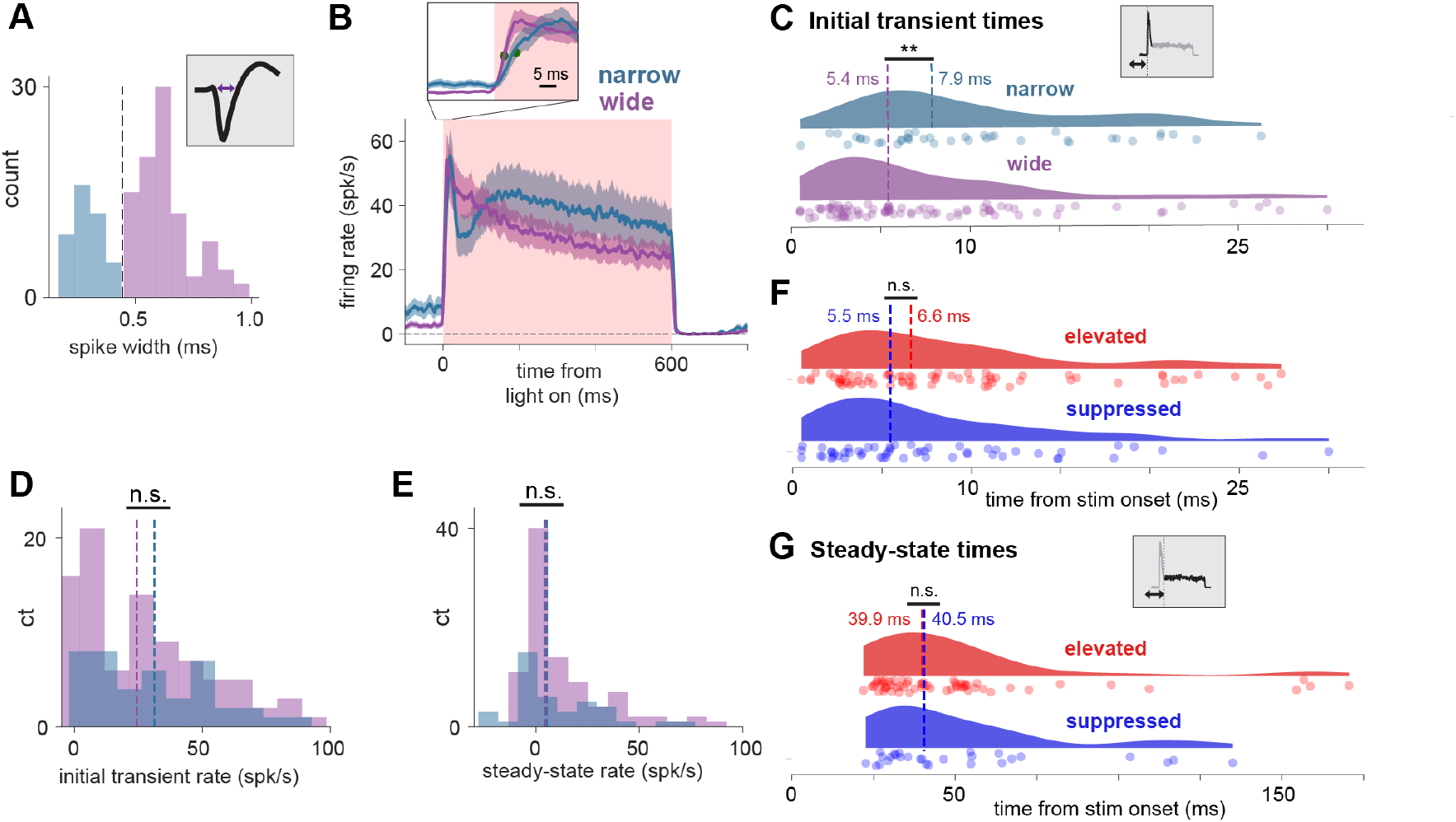
Response dynamics are consistent with steady-state suppression shaped by a recurrent network mechanism. Wide-waveform (excitatory) units have slightly earlier onset latencies than narrow (inhibitory) units, but other quantities do not differ across neural populations. **(A)** Wide/narrow sorting approach. Bimodal widths (classification threshold: 0.445 ms). **(B)** Average traces for narrow- and wide-waveform units (Methods). Inset: enlarged view of laser onset, highlighting latency difference between narrow and wide units. Green markers: time to half peak. **(C)** Onset latencies slightly shorter for wide-waveform units. **(D)** Peak firing rate does not differ between wide and narrow units (medians, wide: 24.5 spk/s, narrow: 31.2 spk/s). **(E)** Steady-state firing rate does not differ (medians, wide: 4.0 spk/s, narrow: 5.1 spk/s). **(F)** Onset latencies do not differ for elevated and suppressed populations. Conventions as in C. **(G)** Time to steady state does not differ for elevated and suppressed populations. See also Fig. S5.

We classified cells into putative excitatory and inhibitory classes by waveform (Fig. 5A). We have previously confirmed^13^ with *in vivo* pharmacology that narrow-waveform cells are inhibitory interneurons, likely PV-positive fast-spiking cells, while wide-waveform cells are primarily excitatory neurons. We saw here that wide-waveform (largely excitatory) neurons have a slightly faster onset latency than narrow-waveform inhibitory cells, faster by approximately 2.5 ms (Fig. 5B, inset, 5C; narrow latency 7.9 ms, wide latency 5.4 ms, difference 2.5 ms, Mann-Whitney U = 1256.0, p < 0.01; onset latencies computed via curve-fitting to rising phases, see Fig. S5A for details).

If subgroups of excitatory and inhibitory cells composed the suppressed and elevated populations, we might expect to see differences in the dynamics of elevated and suppressed cells. But we found no significant differences in onset time or time to steady state for elevated and suppressed neurons (Fig. 5F,G, onset time Mann-Whitney U = 1741.0, p = 0.17, time to steady state, Mann-Whitney U = 801.0, p = 0.32). This was also true when restricting the analysis to only wide-waveform cells (Fig. S5C,D). Another possibility could have been that the neurons with suppressed steady-state responses were cells that did not express opsin. But the similar onset latencies of the elevated and suppressed cells (Fig. 5F) excludes that possibility, and provides further support to the idea that instead a balanced-state recurrent network explains the suppression.

Beyond the differences in onset latency, we found other response dynamics were not different between excitatory and inhibitory cells. Consistent with a recurrent network with strongly coupled excitation and inhibition, we found that both excitatory and inhibitory cell populations increase their average firing rate when excitatory cells are stimulated (wide mean Δ: 14.54 spk/s, t = 5.52, df = 93, p < 0.001, narrow mean Δ: 13.53 spk/s, t = 3.07, df = 41, p < 0.01). That is, both excitatory and inhibitory populations contain elevated and suppressed neurons, though elevated cells dominate both averages (Fig. 5E). Further, the initial transient and steady-state firing rate medians were not detectably different between inhibitory and excitatory cells (transient: Mann-Whitney U = 1816.0, p = 0.23, steady-state Mann-Whitney U = 1866.0, p = 0.31, Fig. 5D,E). Also, time to steady state for wide-waveform and narrow-waveform cells did not differ (Fig. S5B), consistent with the idea that the steady-state dynamics emerge from integration of both inhibitory and excitatory inputs. Overall, the response distribution and dynamics we observed in inhibitory and excitatory cells are consistent with a strongly-coupled recurrent network.

### A neuron’s response is only weakly predicted by optogenetic input to that neuron

We used the imaging data to determine if the suppression we observed was explained by variation across cells in optogenetic drive. We found that while indeed there was variability in different cells’ responses, there was very little relationship between opsin expression and cells’ firing rate changes. To estimate the optogenetic drive to individual neurons, we measured fluorescence of mRuby2 (fused to stChrimsonR) in donut-shaped regions around each cell’s membrane using two-photon imaging (Fig. 6, Fig. S6). The measured *in* vivo distribution of opsin expression was well-fit by a lognormal distribution after excluding the 16.8% (Wilson score 95% confidence interval: [12.7%, 22.1%]) of cells with low fluorescence (Fig. 6C, see Methods). The *in vivo* estimate of the percentage of non-expressing cells was lower than what we observed with histology, perhaps because we selected FOVs with dense opsin for *in vivo* imaging. At the same time, however, our *in vivo* observations are consistent with past work that finds that AAV transfects adult neurons in a non-uniform way, finding substantial variability in opsin expression from cell to cell^13,31^.

**Figure 6:**
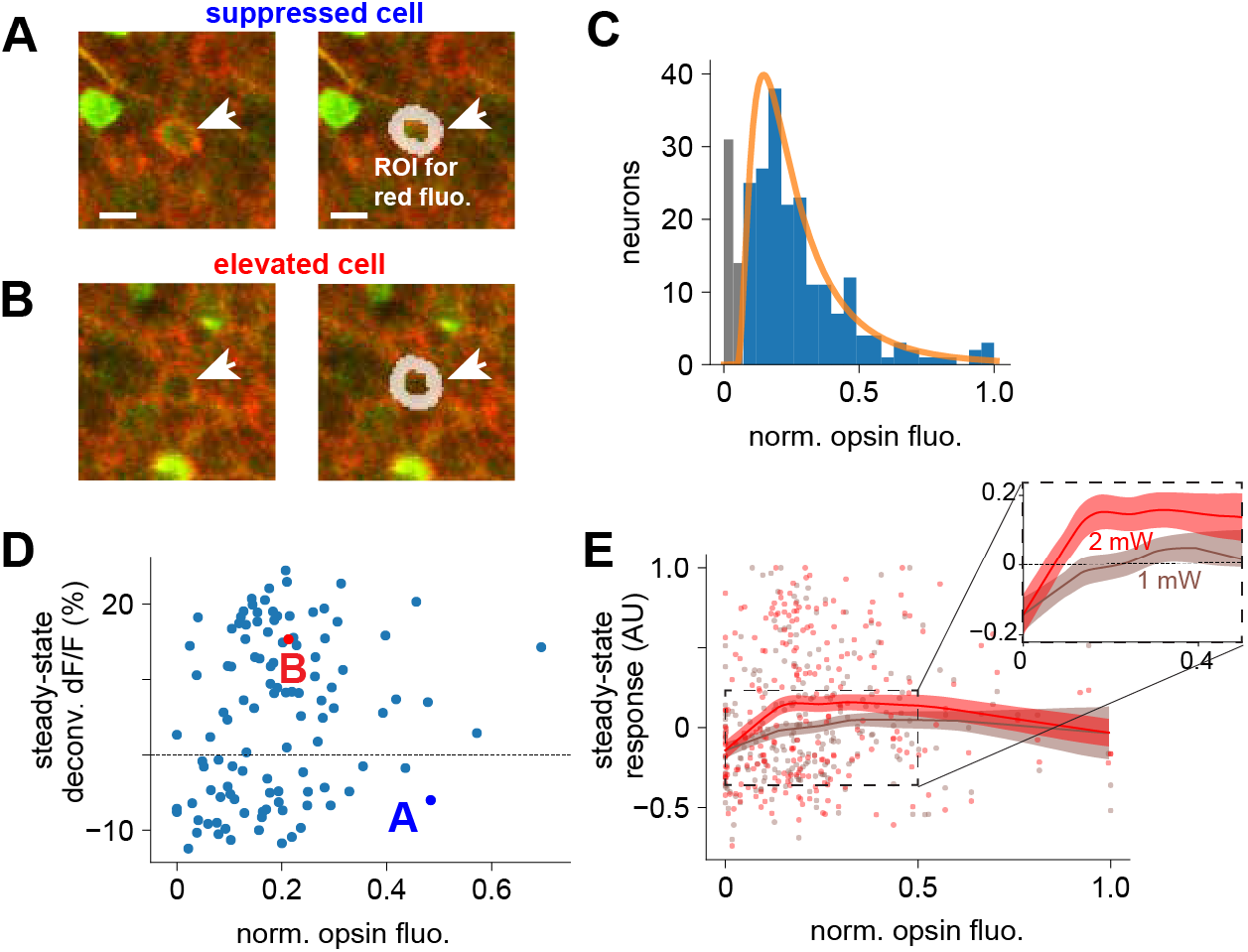
stChrimsonR expression only weakly predicts 2-photon steady-state response. **(A)** Left: Red (stChrimsonR-mRuby2) and green (GCaMP7s) fluorescence of an example cell with a suppressed steady-state response during optogenetic stimulation. Right: donut-shaped region of interest (ROI), inner and outer boundary calculated by shrinking or expanding the cell border (CaImAn, Methods) **(B)** Same as (A), except for an example cell that shows an elevated steady-state response. The suppressed cell shows brighter red fluorescence than the elevated cell, quantified in D. **(C)** Distribution of opsin fluorescence intensity (N=3 animals). Orange: Lognormal fit, excluding non-expressing cells (gray; see Fig. S6). **(D)** Example relationship between opsin expression and response; only a weak relationship is seen. x-axis: red fluorescence in donut-shaped ROIs (n = 113 cells, N=1 animal; 2 mW stimulation power). Example cells are highlighted (colored markers, letters). **(E)** Population data: same as D for N=3 animals (N=244 neurons). Two laser intensities, 1 mW (brown), 2 mW (red). Heavy lines: LOWESS fits; shaded regions: bootstrapped standard error. Slight decline at high values may be due to response saturation or overexpression of opsin in a few cells. Inset: Zoomed view of area indicated by dashed box, cells with the least opsin expression show a slightly smaller response on average than other cells. See also Fig. S6.

We found that the amount of opsin-related fluorescence explained little of the variance in steady-state responses (Fig. 6A-D, Pearson’s r = 0.21, df = 106, p < 0.05, high fluorescence neurons excluded [> 0.5]; Fig. 6D-E, population: at 1 mW: Pearson’s r = 0.18, df = 219, p < 0.01, at 2 mW: Pearson’s r = 0.17, df = 219, p < 0.01).

This striking decoupling effect — that the amount of opsin input barely predicts how cells’ firing rates are modulated by stimulation — suggests that a given cell’s response may not be dictated by input to that cell, but instead by recurrent inputs.

Notably, both high-expressing cells and low-expressing cells showed little relationship between opsin expression and response (Fig. 6E, red and gray lines). This supports the idea that the decoupling is not due to cell-autonomous intrinsic properties but indeed due to recurrent network inputs. To further test this, we measured the variability of neural responses as a function of stimulation intensity. If a neuron’s response were in fact controlled primarily by its opsin expression (the optogenetic input to that neuron) and not network input, increasing the input intensity should keep the variance in response the same, or reduce it, because the fixed opsin level is the principal source of response drive (Fig. S2K). Or, if response was dictated by opsin level, increasing intensity might produce a bimodal response distribution as the optogenetically-driven neurons separate from non-expressing neurons (Fig. S2L). We found support for none of these possibilities. Instead, the response pattern increased in variance as stimulation grew stronger (Fig. S2M-O), supporting the idea that it was network input, not opsin level, that controlled cells’ responses.

We next turned to simulations, fit to our data and building on the recent theoretical advances of Sanzeni et al. (in press), to more completely characterize recurrent network influences on neurons’ responses.

### Input from the recurrent network dominates responses, as explained by a balanced-state model

Thus far, a moderately- or strongly-coupled balanced-state network seems consistent with both the response distributions and dynamics we observe. Indeed, recent theoretical work in rate-based models^22^ has shown that this kind of heterogeneous network response (“reshuffling”) occurs in strongly-coupled cortical networks. To understand if our experimental data could be explained by this reshuffling mechanism, we examined recurrent network models with features reflecting our data, and determined which features of the recurrent network models were important to explain the suppression.

We simulated conductance-based spiking neural networks, varying network connectivity and opsin drive across neurons in these models, and measured network responses to excitatory cell stimulation (Fig. 7A,B).

**Figure 7:**
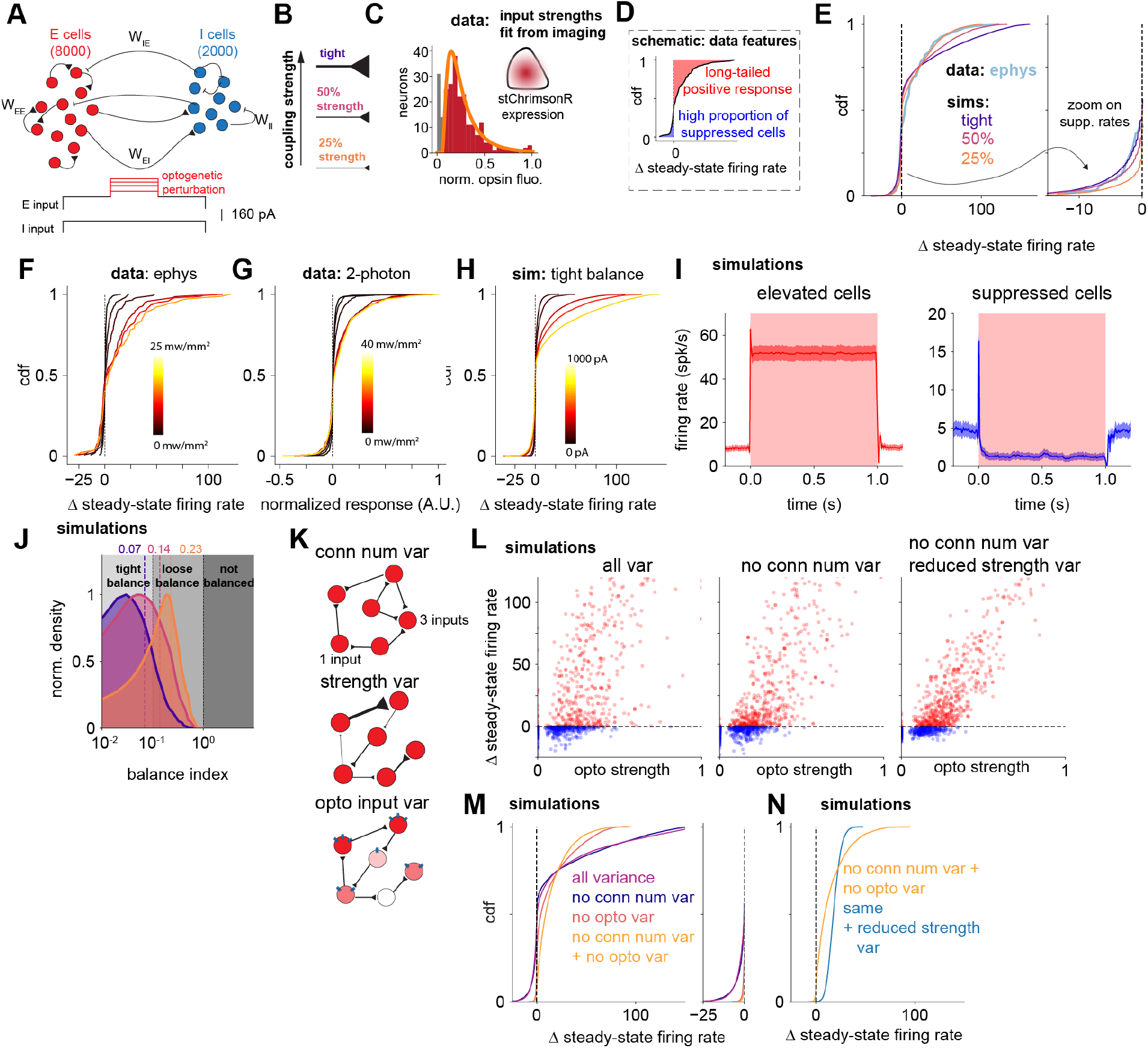
Strongly-coupled recurrent neural network model with heterogeneous connectivity describes the data. **(A,B)** Simulation design: (A) conductance-based spiking network model with 8000 excitatory cells and 2000 inhibitory cells. (B) Network mean recurrent strength is varied to measure effects on neural responses. **(C)** Optogenetic input strengths sampled from a lognormal distribution fit from *in vivo* 2p measurements (Fig. 6C). **(D)** Schematic of data features simulations describe. **(E)** The tightly-balanced model fits the long tail of excitation and the proportion of suppressed neurons. **(F-H)** Responses to stimulation during (F) electrophysiology, (G) 2-photon experiments, and (H) simulations. **(I)** Left, Mean timecourse, elevated cells, strongest recurrent network. Right, same: but for suppressed cells. **(J)** Balance index (Ahmadian and Miller, 2021) of the 3 networks (B). The strongest-coupled network (purple) has a median index in the tight balance regime. **(K)** Schematic of types of input variability. Variation in input can arise from variation across cells in number of recurrent inputs, strength of recurrent inputs, or optogenetic input strength. **(L)** Simulated neural responses to optogenetic input, with (left, same as Fig. 7F) and without (center) variability in number of recurrent inputs. Relationship between optogenetic input and response strengthens when variance sources are removed (R^2^ original = 0.50, R^2^ reduced conn. Num. var. = 0.66; R^2^ reduced conn num and reduced strength var. = 0.77) **(M)** Steady-state firing rate distributions when input variability components are removed. Purple: network with parameters as in panels E (tight),H,I,L. Right: Same data, zoom to the suppressed portion of the distribution. Some suppression exists if either source of variance is removed, but suppression nearly abolished when both sources of variance are removed. **(N)** Finally, reducing variance in synaptic weights (by a factor of 10) nearly removes response variability and suppression. See also Figs. S7-8.

Each simulation consisted of two sparsely connected populations of conductance-based spiking neurons, one excitatory (80%) and one inhibitory population (20%). For each set of network parameters, we adjusted a background input current to either excitatory or inhibitory neurons to hold the spontaneous firing rate of the neurons at a value (∼5.4 spk/s) consistent with the data (Fig. S8F). We drew the opsin input strength for each neuron from a distribution fit to the imaging data, and scaled that distribution until the 75^th^ percentile of the network response matched the electrophysiology data (lognormal distribution, with 16.8% nonexpressing, Fig. 7C). We also ran these simulations using the percentage of nonexpressing neurons as estimated from the histology data (41%) and found no qualitative differences (Fig. S8I-M).

We first manipulated the mean connectivity strength of recurrent connections. We constructed three different models, varying the average strength of recurrent coupling in each (schematic, Fig. 7B). The “tightly balanced” network had the strongest recurrent coupling, and we scaled down the synaptic weights by a factor of 2 or 4 to create more weakly-coupled “50%” and “25% strength” network simulations. We confirmed that each of these simulations showed paradoxical suppression of inhibitory cells, a sign of strong recurrent coupling within the excitatory network and the ISN regime, as observed in visual cortex^11,13,34^. We stimulated the inhibitory cells in each network and found, as expected, paradoxical suppression (Fig. S7A,B).

Recurrent excitatory-inhibitory networks can be tightly or loosely balanced^10^, depending on the total amount of recurrent excitatory and inhibitory input to network neurons. To classify the networks, we calculated the balance index, a ratio that measures how completely inhibitory input cancels out the excitatory input for each neuron in the networks^10^ (see Methods). We found that all three networks we constructed are balanced, as expected due to their irregular spontaneous activity (balance index << 1), and the networks span a range from loose to tight balance (Fig. 7J).

### The model replicates the long tail of positive responses, suppressed responses, and dynamics

Two characteristic features of the response data we observed are the long-tailed positive response and the substantial proportion of suppressed cells (Fig. 7D). All three simulated networks showed a long tail of elevated responses as in the data, with many neurons showing increases in firing rate to stimulation, and a few showing large increases (Fig. 7H). However, the amount of suppression depended on recurrent coupling strength. Increasing the total excitatory and inhibitory recurrent input by varying the mean coupling strength leads to more suppressed neurons when other network parameters are held constant (Fig. 7E). The network that best fit the fraction of suppressed cells we observed was the most strongly-coupled network, which was just inside the tight balance regime (Fig. 7E,J; suppression sensitivity to baseline rate and coupling strength characterized in Fig. S8F-H). Additional model components could lead to similar results with networks of higher or lower balance index estimates. For instance, adding structured connectivity may reduce the required coupling strength^22^. However, our data underline that the recurrent coupling should be strong enough so that when input arrives to a population of neurons, many neurons’ responses are substantially controlled by their recurrent input.

Excitatory and inhibitory cells’ response distributions were similar in the model, as also seen as in the data (Fig. S7E,F).

Finally, to further demonstrate how well the model could reproduce the features of the observed response distributions, we simulated responses while parametrically increasing the strength of the input, and compared the results to the electrophysiological and 2-photon responses to increasingly strong experimental optogenetic stimulation. The shapes of the response distributions in the most tightly coupled model and the data were similar (Fig. 7F–H).

Given the ability of a balanced-state model to describe the suppression, we checked if the model dynamics were consistent with the data. We found that model responses were qualitatively similar (Fig. 7I) to the timecourses of responses seen in the data (Figs. 2–3). Excitatory cells first showed a brief, positive transient response before the network settled into a new steady state, with some cells excited and some suppressed. The initial positive transient in suppressed cells is a key observation, as it suggests a network mechanism where input initially excites many excitatory neurons, but later recurrent inputs lead to suppression in many of the same neurons. A second similar feature of the dynamics in model and data is the offset dynamics in both elevated and suppressed cells: after stimulation ends, both show a slight suppression before returning to baseline. Finally, excitatory cells have earlier onset times, indicating that E cells were directly stimulated and I cells were recruited just a few milliseconds later (Fig. S7D), before both populations then evolved to a new steady state.

One feature of the dynamics seen in the data but not the model is that for high stimulation powers there is a slight decay during the tonic or steady-state period (Fig. 3I). However, this decay effect is not seen at lower stimulation intensities, suggesting it arises from known opsin dynamics (inactivation at high light power, e.g.^35^) or other known biophysical, non-network, effects like spike-rate adaptation.

In sum, this model recapitulates many of the features of our observations, suggesting that a excitatory-inhibitory mechanism with strong and variable recurrent coupling explains how V1 neurons respond to input.

### Variability in recurrent input creates different responses in different cells, and explains the decoupling of a neuron’s response from its optogenetic input

In our data, we saw strikingly little correlation between opsin expression and neural response to stimulation (Fig. 6), suggesting recurrent input strongly governs the response. In fact, the tightly-coupled model showed the same pattern of responses (Fig. 7L, left).

We therefore asked which sources of variability were important to explain why neural responses were weakly related to optogenetic input. To do this we varied sources of input variability in the model (Fig. 7K). First, we reduced variability in either the number or strength of recurrent inputs and found that this created a stronger correlation between optogenetic input strength and response (Fig. 7L, middle and right) which made the model a worse fit for the data (Fig. 6). The original relationship was not recovered by increasing the recurrent strength of the network (Fig. 7 Supp 3), implying recurrent connection variability was required to produce this effect and higher recurrent strength could not substitute for it. Next, we asked whether variability in opsin input across cells was also essential to explain the suppression we observed. Removing the variance in optogenetic input across cells (so that each excitatory cell with opsin received the same input) significantly reduced the number of suppressed neurons and also produced a worse fit to the data (Fig. 7M; right inset highlights suppressed neurons).

If input variability was the primary source of variability in neural responses, then removing variability from both kinds of input — both optogenetic input variability and recurrent input variability — should substantially reduce the amount of suppression observed. This is what we found. Removing or reducing both types of variability produced a set of neural responses clustered tightly around the mean response (Fig. 7N) with no suppressed neurons. Thus, both variability in recurrent input and in optogenetic input are required to explain the data.

Together these results show that both optogenetic input variability and recurrent connection variability help create the variability in different neurons’ responses. Each neuron’s firing rate is affected not just by the optogenetic input that particular cell receives, but also by recurrent input received from other neurons, and the other neurons themselves receive different amounts of optogenetic and recurrent input. When optogenetic input is delivered, the whole network changes state to a new set of firing rates, and each neuron’s new firing rates are only weakly related to the optogenetic input to that neuron. Thus, the recurrent network explains the unexpected decoupling of optogenetic input strength from neural response strength that we observed experimentally (Fig. 6).

### A balanced-state network model with connection variability also explains expected responses to single cell stimulation

Past work has found that stimulating a single cell in visual cortex leads to mean suppression in the surrounding population^20^ Our results seem initially to contradict this finding, because our data and simulations both find a net positive response across the population when we stimulate many excitatory cells.

To determine if the effects of single-cell stimulation could also be explained by the balanced-state simulation that describes our data, we performed simulations of single cell stimulation in the same tightly-coupled network (Fig. 7A,B), measuring the response of the non-stimulated population (Fig 8A,B). We found that, while single cell stimulation produced a range of individual cell responses (i.e. reshuffling, Fig. 8C), the mean response was not negative, but instead close to zero (Fig. 8C,D; mean firing rate 95% CI [-0.006, 0.021], t (9998) for nonzero mean = 1.13, p = 0.26). The excitatory cells that received a direct connection from the stimulated cell (feedforward, FF, cells, n=107 neurons) had an elevated response. Those that received a connection from the inhibitory cells which received a monosynaptic input from the directly connected E cells had a very slightly suppressed response (i.e. E-I-E connections, or feedback (FB) cells, n=4380 neurons; Fig. 8E). The small set of strongly excited cells average with the large number of weakly suppressed cells to lead to a mean response near zero (Fig. 8F,G).

**Figure 8:**
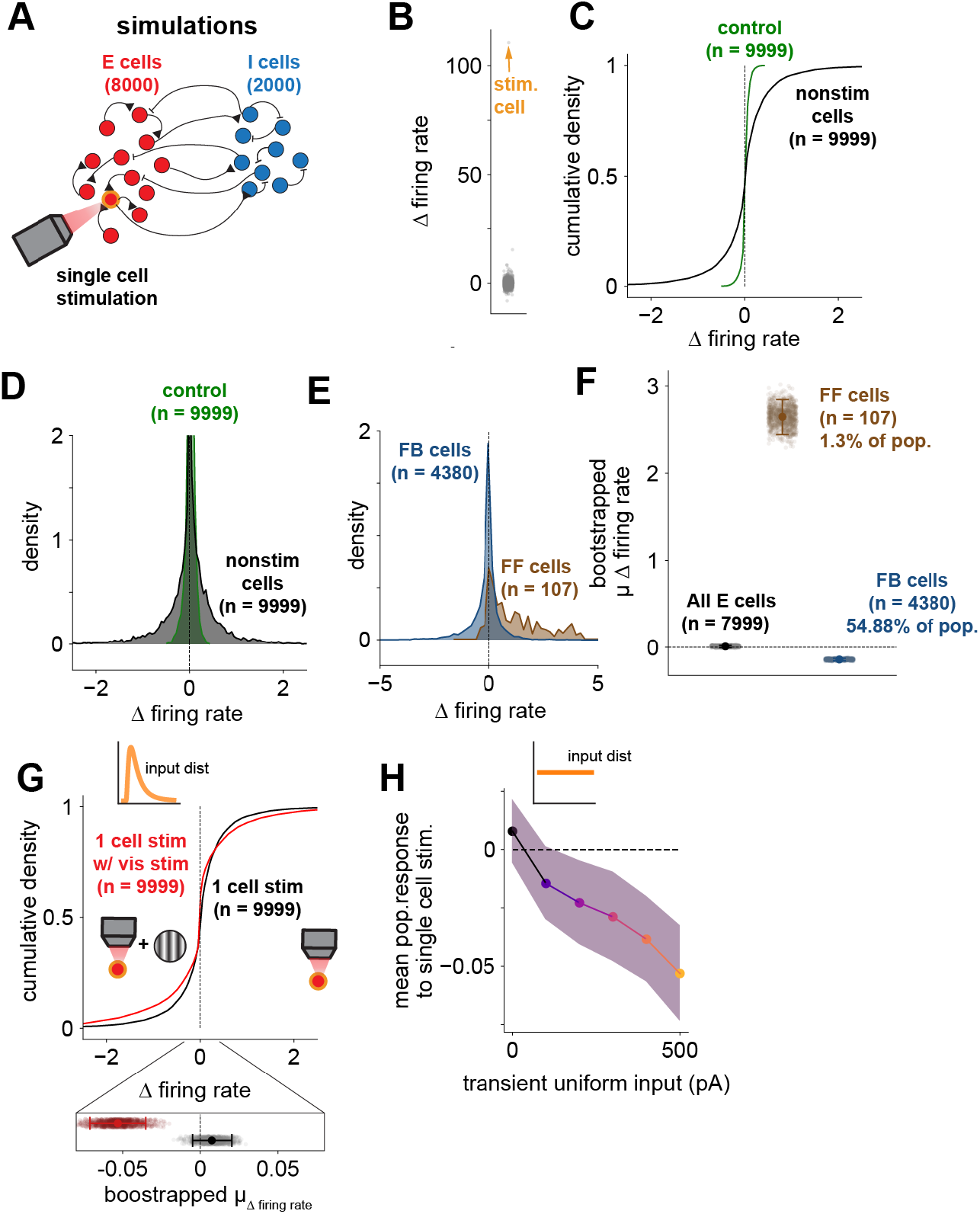
Single cell stimulation produces elevated firing rates in a small subset of cells, but widespread weak suppression across the population. **(A)** Simulation schematic: one cell stimulated (‘tight’ network, Fig. 7B,E,H). **(B)** Single cell stimulation weakly modulates other cells. **(C)** Single cell stimulation reshuffles the distribution of responses; individual neurons change response (black, note variance of distribution), mean/median remain near zero. **(D)** Densities, same data as (C). **(E)** Defining cells by their connectivity to/from the stimulated cell (direct input from stimulated cell, FF, brown; input from an inhibitory cell receiving FF input, blue) reveals a small number of excited cells. N=1 instantiation of network (weight choice). **(F)** Means of E across many instantiations. Black: full population of E cells, Brown, blue: same conventions as E. Error bars: SEM. **(G)** Simulated visual input during stimulation leads to mean suppression. Red: lognormally distributed input + single cell stimulation, black: single cell stimulation alone. Red mean is negative. **(H)** Mean suppression increases with stronger input.

Single-cell stimulation in the model we fit, therefore, could not account for the mean suppression observed in previous studies. We hypothesized that this difference could be due to difference in the activation state of the network. Chettih and Harvey (2019) stimulated during visual input, while here we delivered optogenetic input during spontaneous activity. Such effects can be seen in balanced-state models: Sanzeni et al. (in press) found that increasing the firing rate of a similar network to our model reduced the mean response of the network. Further, in models of visual cortex with subnetwork connectivity (e.g. higher connectivity between neurons with similar orientation tuning^36^, it has also been shown that visual input can shift the network response to be more negative^37^. Therefore, to test whether additional network drive could reproduce mean suppressive responses, we simulated single-cell stimulation paired with an input that mimics visual drive (Fig. 8G; Methods). Indeed, this shifted the mean population response negative (Fig. 8G; firing rate change: mean: -0.05; 95% CI: [-0.07, -0.04]). This effect is quantitatively dependent on the strength of the simulated visual input: as simulated visual input grows stronger, the more negative the mean response to optogenetic input becomes (Fig. 8H).

Thus, a strongly-coupled balanced state model is consistent with not just our data, but with past results on single-cell stimulation. Strong mean connectivity, as well as variability in recurrent connectivity, shape the responses of the network.

## Discussion

We found robust suppression in visual cortex in response to direct optogenetic drive to excitatory neurons, with intermixed elevated and suppressed neurons. This salt-and-pepper distribution of responses resembles what is observed during visual input, and arises without input to inhibitory neurons. The firing rate distributions and response dynamics suggest a network mechanism for the observed suppression: that recurrent input variability, combined with external input variability, decouples the optogenetic input strength from the firing rate response in individual cells. This yields a weak correlation between input and response (Fig. 6E), so that a high level of opsin in a cell does not necessarily mean that cell fires strongly in response to stimulation. This recurrent network mechanism seems likely to create variability in visual responses as well (Fig. 1), because these recurrent connections are present in the cortical network for all kinds of input, and so shape responses to visual input also.

Intuitively, the network mechanism that creates the salt-and-pepper excitation and suppression is that external inputs first elevate the firing rates of excitatory cells (Fig. 5A-C), some more than others. That activation excites inhibitory cells, also some more than others. The result is the network settles into a new steady state (Fig. 5E–G) with a very broad distribution of excitatory cell firing rate changes (Figs. 6,7). Our measurements, showing a long tail of excited responses (Fig. 2I), a substantial number of suppressed cells, and response dynamics with initial transients followed by steady-state excitation and suppression (Fig. 3I,J), all confirm that recurrent inputs can explain the response patterns we see.

The salt-and-pepper pattern of responses varies gradually over space, with suppressed cells becoming a larger proportion of neurons with distance from the stimulation site (Fig. 3,4). The salt-and-pepper distribution of responses we observe is therefore overlaid on top of the global trends we observed in widefield imaging. This global suppression, in a concentric surround region similar to surround suppression during vision (e.g.^38^), is driven by direct excitatory inputs, suggesting that visual surround suppression is not inherited from other regions but also arises from recurrent interactions.

### The role of inhibition and suppression in the cortex: sharpening or high-dimensional pattern modification?

In principle, one role of suppression in the cortex could be to sharpen responses to input via attenuating responses in non-driven cells. The finding of distance-dependent suppression in our widefield data (Fig. 4) implies exactly this conclusion. Pioneering work using single-cell stimulation^20^ also found the same sort of suppression in non-stimulated neurons. They showed that suppression falls off with distance by averaging across recorded neurons. (Note that this is true across tuning properties: while like-tuned cells in Chettih and Harvey (2019) show less suppression than other neurons, the average effect in like-tuned cells is still suppression.)

Our data and model extend this to show that sharpening is not the only, or likely even the primary, effect of cortical suppression. Using individual cells’ responses with 2p imaging and electrophysiology combined with simulations, we demonstrate that cortical stimulation generates large response variability even in cells directly receiving input. That statement has significant consequences for how the cortex transforms its input — it is not just that recurrent connectivity sharpens responses, but it can create much more complex and high-dimensional transformations^21^. Such transformations are central to neural coding and how neural codes are created from input.

### Variability in recurrent connectivity in the cortex: experimental evidence

We find that variability in connection strength between L2/3 excitatory neurons is necessary to create the heterogeneous responses to input we, and others^39^, observe. Several observations suggest that the brain has recurrent variability at least as large, and possibly larger, than we use in the simulations. First, electrophysiological studies often find a long tail of synaptic strengths between pairs of neurons, with a few very large connections^40–42^. The variance of individual synaptic weights may be lower^43^, with the larger connection strengths due to multiple synaptic contacts between neurons (though see^44^ for evidence of long-tailed synaptic bouton sizes.) If there is a long tail in synaptic connection strengths, this would still support our finding of high recurrent variance, as it would increase the recurrent variability even beyond the weight distributions we used, which are truncated Gaussians with mean and variance equal. Second, we used a connection sparsity of 2%. We set the number of inputs a cell received from the recurrent network according to a binomial sum, with fixed connection probability between neurons. Connection probability in the brain may be higher, as for example paired recording studies have found connection probabilities of 10% or higher^41,42^. And higher connection probability will produce greater variance in net input into different cells, as binomially-distributed sums have a larger variance as connection probability increases (in the 0–50% range). Finally, patterned or subnetwork-specific connections, which we did not include, would also only increase variance, though specific connections seem to have just a moderate effect on connection probability — shared tuning changes the connection probability from 10–20% on average to 30–50% for like-tuned neurons, in some cases^36^. Taken together, the substantial recurrent variability that explains our data is consistent with experimental measurements of recurrent connection variability.

### Strong balance, loose balance: implications for models that describe cortical networks

We find that a two-population excitatory-inhibitory model is sufficient to explain the data we observe. *A priori*, it could have been that a model with multiple inhibitory subtypes^15,45^ would be needed to reproduce the dynamics and population statistics we saw. Recent work has argued for particular roles for cortical inhibitory subtypes: that parvalbumin-positive (PV) neurons are the primary class providing inhibition stabilization^13,46^, while somatostatin-positive (SOM) cells are involved in gain control^46^. These separate roles are still consistent with our findings. PV cells are likely to be the primary inhibitory cell class in our data and model, as PV cells are the narrow-waveform cells that we identify in electrophysiology^13^ (see Fig. 5). Those cells show dynamic and response firing rate changes expected for the inhibitory population in an E-I model (slightly delayed onset latency, similar distribution of firing rate change as E cells). It is also plausible that stimulating cortical excitatory cells as we did does not cause gain to vary, so that a separate gain role of SOM neurons was not evident in our experiments. Thus, a two-population inhibitory model (with PV cells likely making up a large part of the I population in the model) is sufficient to explain our data.

In addition to supporting the idea that recurrent connections between neurons have substantial variability, our results also confirm that the mean V1 recurrent connectivity is strong — i.e. V1 operates as an inhibitory-stabilized network, meaning that the excitatory network is unstable if inhibition could be frozen^11,13,14,22^. Within the class of balanced networks, two sorts of balance have been distinguished: “loose” and “tight” balance^10^. The best network in our results (Fig. 7) is on the border of the tight- and loose-balance regimes, with individual cells falling in either the tight or loose-balance regimes. A network near the transition from loose to tight balance is broadly consistent with past experimental data (^22^, reviewed in ref. ^10^) which do not suggest a very tightly-balanced regime for the cortex (Fig. 7). Recent work has shown that adding structured (tuned subclass) connectivity allows substantial recurrent effects with looser balance^22^, further supporting the idea that our data support loose or moderate balance.

The mechanism we find for suppression is strikingly different than paradoxical suppression in an ISN when inhibitory cells are stimulated^12–14^. In both cases, suppression is paradoxical: here we excite excitatory cells and see suppression of excitatory cells, and in an ISN, exciting inhibitory cells causes suppression in inhibitory cells. But in paradoxical inhibitory suppression, the *mean* firing rate of the inhibitory population decreases^14^. Here with excitatory cell stimulation, the mean firing rate change is non-paradoxical, as excitatory cell average rates *increase*. It is the substantial variability or heterogeneity of recurrent connections in combination with variability of input that causes many cells to be suppressed as others increase their firing. However, both types of paradoxical suppression, when excitatory or inhibitory cells are stimulated, are only present when the network operates as an ISN – that is, both effects happen in a network with strong average recurrent coupling^13,22^. The observed paradoxical suppression of excitatory cells, however, requires variability around that strong average recurrent coupling.

### Future: subnetworks, computation, and interactions between areas

These results could be extended in a few ways. First, here we did not consider how subnetwork connectivity between excitatory neurons in the cortex might influence the effects. Ko and colleagues (2011) showed approximately a 2–3 fold increase in probability of connection between V1 excitatory neurons that had similar tuning (orientation or direction) compared to those with dissimilar tuning. Adding subnetwork connectivity would not qualitatively change our conclusions: that suppression results from recurrent influences, and that it depends on variability of connectivity within the network. However, future work stimulating within or across subnetworks might change the fraction of cells suppressed, given that input patterns would drive neuron populations with somewhat more or less connectivity with each other and the rest of the network. Cell-specific two-photon holographic stimulation^16,47–49^ seems well-placed to study how patterned activity in one subnetwork affects activity in another subnetwork.

While local collaterals probably contribute the majority of recurrent cortical input, meaning nearby neurons influence each other via direct synapses, it is possible that long-range, inter-areal, connections could contribute to the experimental results we observe. Estimates of connectivity falloff show most connections to a given neuron come from local neurons^6,7^. But in principle, cells in other areas could form part of the recurrent population. This could happen for example if projections from V1 to the thalamus recruited neurons there which connect back to the cortex. However, our widefield imaging data (Fig. 4) shows that the suppression peaks a few hundred microns from the stimulation site, suggesting relatively local influence. Therefore, it seems likely that the recurrent connections in the simulations primarily reflect local connections within V1 to nearby neurons.

## Conclusion

Here we find paradoxical suppression of excitatory cells in the cortex when excitatory cells are stimulated. These results suggest that a primary purpose of recurrent connectivity in visual cortex is to change the steady-state firing rate of network neurons, beyond just how inputs are transformed by feedforward connections. Our results are a step forward in explaining how cortical networks change their firing in response to different patterns of input — a fundamental building block of neuronal computation.

## Acknowledgements

We thank Kaya Matson for help with RNAscope, Aanika Kashyap for histological analysis.and Nicolas Brunel, Alessandro Sanzeni, and Ken Miller for helpful comments on the manuscript and/or discussion. This work was funded by the National Institutes of Health (BRAIN U01NS108683 and intramural support ZIAMH002956). This work utilized the computational resources of the NIH HPC Biowulf cluster (http://hpc.nih.gov).

## Author contributions

Electrophysiology data was collected by ZZ. 2-photon data was collected by PKL, AJL, and JFO. Widefield data was collected by JFO. Histological data collection and analysis was performed by ZZ and HCG. JFO and MHH designed and implemented simulations. JFO, MHH, and ZZ curated and analyzed data. JFO and MHH wrote and edited the manuscript.

## STAR Methods

### Key resources table

| Reagent or resource | Source | Identifier | Additional information |
| --- | --- | --- | --- |
| *Chemicals, peptides, and recombinant proteins* |  |  |  |
| Tamoxifen | Sigma-Aldrich | T5648-5G |  |
| *Experimental models: organisms/strains* |  |  |  |
| Ai148 | The Jackson Laboratory | RRID:IMSR_JAX:030328 |  |
| Ai162 | The Jackson Laboratory | RRID:IMSR_JAX:022731 |  |
| Cux2-CreERT2 | MMRRC | RRID:MMRRC_032779-MU |  |
| Emx1-cre | The Jackson Laboratory | RRID:IMSR_JAX:005628 |  |
| *Bacterial and virus strains* |  |  |  |
| AAV9-hSyn-jGCaMP7s-WPRE | Addgene | RRID:Addgene_104487 |  |
| AAV9-Syn-DIO-stChrimsonR-mRuby | Addgene | RRID:Addgene_105448 |  |
| AAV9-CamKIIa-ChrimsonR-mScarlet-KV2.1 | Addgene | RRID:Addgene_124651 |  |
| *Software and algorithms* |  |  |  |
| Mworks | The Mworks Project |  | mworks.github.io |
| *Other* |  |  |  |
| C and B Metabond | Parkell | S380 |  |
| Kwik-sil | World Precision Instruments | KWIK-SIL |  |

### Resource Availability

- *Lead Contact* Additional information and requests for resources should be directed to the lead contact, Mark Histed
- *Materials availability* This work did not produce novel reagents.
- *Data and code availability* Data and code will be published in a GitHub repository or on DANDI on acceptance for publication.

### Experimental Model and Subject Details

All procedures were approved by the NIMH Institutional Animal Care and Use Committee (IACUC) and conform to relevant regulatory standards. Emx1-cre animals^30^ of both sexes (N = 14; https://www.jax.org/strain/005628) were used for 2-photon and electrophysiology experiments (5 for electrophysiology, 3 for visual stimulation imaging, 6 for optogenetic stimulation imaging). For widefield imaging experiments, Ai162 (N = 2; https://www.jax.org/strain/031562) and Ai148 (N = 1; https://www.jax.org/strain/030328) animals^50^ were crossed with the Cux2-CreERT2 line^51^ (https://www.mmrrc.org/catalog/sds.php?mmrrc_id=32779), and GCaMP6f or GCaMP6s was induced via tamoxifen injection during adulthood (P22 or later, tamoxifen 2 mg intraperitoneally daily for 3 days). All animals were singly housed on a reversed light cycle. During experiments animals were water scheduled and given occasional water rewards to keep them awake and alert. To ensure animals did not drift into a quiet wakefulness or quiescent state, we monitored animals during data collection to verify they continued to drink the delivered reward.

## Methods Details

### Implants and injections

Details of the headplate and window procedures are described in previous studies^13,52^. Optical glass windows (3 mm diameter) were placed over V1 (center: -3 mm ML, +1.5 mm AP, relative to lambda) for 2p and widefield imaging. Windows were also used before electrophysiology for imaging to localize V1.

For Emx1-Cre animals, 300 nL of AAV9-syn-jGCaMP7s-WPRE (RRID:Addgene_104487) and/or AAV9-Syn-DIO-stChrimsonR-mRuby (RRID:Addgene_105448) were injected 250 µm below the dura (200 nL/min) prior to cementing the cranial window. For Ai148 and 162 animals, AAV9-CamKIIa-stChrimsonR-mRuby2 was generated by cloning the CaMKIIa promoter (RRID:Addgene_120219) into a pAAV backbone containing stChrimsonR-mRuby2 (RRID:Addgene_105447) and packaged into an AAV (Vigene, Inc.). This was injected at the same depth as the hSyn-DIO-stChrimsonR virus, but with 100 nL volume at 100 nL/min.

### Electrophysiology

Electrophysiological methods are described in detail in previous studies^13^, and are summarized here. Animals were head-fixed during recording. Before the first session of electrophysiology, the animal’s cranial window was removed and the craniotomy was flushed with saline. Between recording sessions, the craniotomy was covered using Kwik-Sil polymer (WPI, Inc.). A fiber optic cannula (400 µm diameter, 0.39 NA, Thorlabs) was placed to center light output at the center of stChrimsonR expression. For light intensity calculations, spot area was defined as the area inside the 50% contour of light spot intensity on the cortex, measured with a camera by imaging the spot on the brain surface. 1–2% agarose (Type IIIA, Sigma) was placed over the dura at the start of each session, and an array of four electrodes (4 probes, 32 sites in total, part #A4x8-5mm-100-400-177-A32, NeuroNexus, Inc.) were lowered into the cortex using a micromanipulator (Sutter MPC-200). One probe was placed at the center of the light spot.

Probes were advanced 600–1000 µm below the point in which the first probe touched the dura. Probes were not moved for 1 hour prior to recording, as we found this improved recording stability. Recording data was sampled at 30 kHz (Cerebus, Blackrock Microsystems.)

Optogenetic stimulation was performed with randomly interleaved stimulation light pulses with several intensities over the range 0.2 mW/mm^2^ to 15 mW/mm^2^. Stimulation pulses were 600 ms long and delivered with a 4 s period.

For spike recordings, waveforms (bandpass filtered, 750 Hz – 7.5 kHz) were digitized and saved by storing a short data section around points where amplitude exceeded 3 times the RMS noise on that channel. Single units were identified (OfflineSorter, Plexon, Inc) based on clusters in the waveform PCA that were separate from noise and other clusters, had unimodal spike width distributions, and inter-spike intervals consistent with cortical neuron absolute and relative refractory periods. A single-unit score was assigned to each unit manually based on these factors ^13,23^. To compare these populations quantitatively, we calculated SNR for both single and multiunits^53,54^. Median SNR for single units was larger than median SNR for multiunits (SU: 3.32, MU: 2.26; Fig. S2C), consistent with prior reports^13,53,55^.

### Histology

Following completion of electrophysiology experiments, mice were anesthetized with isoflurane and injected intraperitoneally with pentobarbital sodium (150 mg/kg), and perfused transcardially with cold (4°C) PBS followed by cold 4% paraformaldehyde. Brains were extracted and fixed in 4% paraformaldehyde for 6–12 hr and then cryoprotected in a 30% (% w/v) sucrose solution in PBS until they sank. Tissue was cryosectioned at 10µm and mounted on charged slides. Fluorescence in situ hybridization was done using RNAscope’s Multiplex Fluorescent Assay v2^56^. Inhibitory neurons were labeled with a VGAT probe (Slc32a1, #319191-C2; Alexa Fluor 488), excitatory neurons were labeled with a VGLUT1 probe (Slc17a7, #416631-C3; Cy5), and an mRuby2 probe (#487361; Cy3) labeled stChrimsonR-mRuby2 expressing neurons. Slides were coverslipped with DAPI. We imaged slides on a Zeiss LSM780 confocal microscope with a 40x oil immersion objective. We imaged each fluorophore separately with a single excitation laser, and collected all three emission channels. To compensate for bleedthrough where the other two fluorophores might be weakly excited by a laser selected for another fluorophore, we subtracted a scaled version of the primary emission channel image for each non-selected fluorophore from the primary channel for the selected fluorophore. Five representative areas were quantified independently by two observers.

### Visual stimulation

Visual stimuli were presented using MWorks (https://mworks.github.io/). Grating stimuli (sinusoidal contrast variation, 0.1 cyc/deg, orientation = 0 deg) were masked with a circular raised-cosine envelope (15 deg FWHM). Visual stimuli were displayed on an LCD display, with center positioned 0-10 degrees of visual angle temporal to the central meridian. Oriented noise stimuli were generated by filtering white noise pixel arrays (each pixel drawn independently from a uniform distribution) with a spatial band-pass filter (peak orientation = 0 deg, orientation bandwidth = 10 deg, peak spatial frequency = 0.05, frequency bandwidth = 0.05). Frames were generated at 60 Hz and the noise pattern was independent from frame to frame^26–28^. Visual stimuli were presented for 3 or 5 seconds, depending on the experiment.

### 2-photon imaging

During 2-photon experiments, animals were awake, water-scheduled, and given periodic water rewards (20% probability per trial, reward once every 30 s on average). If animals stopped licking in response to the rewards, data collection was ended. We imaged GCaMP7s responses (920 nm excitation) with either a galvo-galvo (5 Hz) or resonant scanning (30 Hz) two-photon microscope. stChrimsonR-mRuby2 expression was imaged at 1000 nm. The microscopes used for imaging were built using MIMMS components (https://www.janelia.org/open-science/mimms-21-2020) and other custom components, built in-house or provided by Sutter Inc. A second light path, combined with the 2p stimulation light path before the tube lens using a dichroic, was used to stimulate stChrimsonR using 530nm light (CoolLED, pE-4000). For 200 ms long optogenetic pulses, we measured responses in the first frame after stimulation. For 4 s long optogenetic light pulses (6 s period), we imaged while stimulation was ongoing. To do this, we avoided stimulation artifacts by stimulating only during horizontal flyback (approximate pulse duration 19 µs, off time 44 µs, duty cycle 30%, line rate 8kHz).

### Widefield imaging

For widefield imaging experiments, we used Ai162;Cux2-creERT2 or Ai148;Cux2-creERT2 animals, expressing GCaMP6f or 6s in L2/3 excitatory cells. Animals were head-fixed and awake during widefield imaging experiments. Prior to imaging, a fiber optic cannula was aimed at the center of the focal stChrimsonR expression. Images were collected using a Zeiss microscope (Discovery V12) with a 1.0x objective using excitation light with wavelength centered at 475 nm (Xylis X-Cite XT720L). A Zyla 4.2 sCMOS camera (Oxford Instruments) collected images (100 ms exposure time, approximately 140 ms frame period) with 4-pixel binning. Laser powers were randomly interleaved, with 50 repetitions per laser power. Laser pulses were 600 ms long, and presented with 6 s period.

### Analysis of electrophysiology data

For spike rate plots, spike counts were binned (1 ms bins), and smoothed via LOWESS^57^. To classify units as having elevated or suppressed responses, we measured spike rate over 145– 400 ms after stim onset, relative to baseline (-1020 ms–0 ms relative to stim onset) for 6 mW/mm^2^ stimulation intensity. To classify cells as wide- or narrow-waveform, we used a spike width threshold of 0.445 ms based on the bimodal distribution of waveform widths (Fig. 5B). This threshold is consistent with pharmacological segregation of inhibitory and excitatory cells^13^.

For analysis of onset times, we fit a sigmoid (logistic function) to each cell’s response from 100 ms before to 100ms after laser pulse onset:

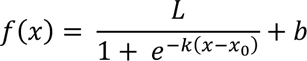

L: upper asymptote, b: lower asymptote, k: slope, x_0_: onset latency. X_0_ was constrained to the range [onset+0.5 ms, onset + 30 ms]. We defined onset latency as x_0_, the time to half-max. To estimate the time to steady-state, the same function was fit to data from 500 ms before and after the laser onset, with the spike rates within a 50 ms window around the initial transient blanked by setting to the baseline firing rate. Each cell’s time to steady-state was computed as the difference between the steady-state onset and the initial onset (difference between the x_0_ parameters of the two fits).

### Analysis of 2-photon data

For short optogenetic stimulation (200 ms pulses) during two-photon imaging, we avoided stimulation light influencing imaging responses by measuring responses in the frame after the stimulus offset. For long pulses (4 s), we stimulated during imaging frames by restricting stimulation to imaging line flyback and intensities we give are the average intensity, corrected for the 30% stimulation duty cycle. Because we found that the LED device we used for stimulation (pE-4000, CoolLED Ltd; specified bandwidth 100 kHz) had some variability in onset/offset for each line, we removed pixels (∼40% of frame) at left and right edges of field of view to ensure no stimulation light could affect images. Image frames were motion corrected using NoRMCorre through CaImAn^58^. Deconvolution was done with OASIS^59^ via CaImAn. To ease interpretability of the deconvolution signals, each neuron’s deconvolved signal was normalized to have the same maximum value as the dF/F of the corresponding fluorescence trace. To separate populations into elevated and suppressed cells, we performed a one-sample t-test (α = 0.05, different from zero, two-tailed) on the deconvolved dF/F during the stimulation period (long pulses) or the frame just after the stimulation period (short pulses). For the short pulses, we used the frame just after stimulation to estimate responses for each neuron per trial. For the long pulses, we averaged data within the period 750 ms after stimulus onset to the stimulus offset in order to capture the steady-state response. For visual response data, data were preprocessed in the same manner as the short optogenetic stimulation experiments. We averaged steady-state responses from 750 ms after stimulus onset to stimulus offset.

For spatial analyses, we used the spatstat package^60^ in R (ver. 4.2.3). For each individual animal, we tested the spatial distributions of elevated and suppressed responses against an inhomogeneous Poisson process model using the *Linhom* and *Lcross.inhom* functions. We used an inhomogeneous process as signal properties (e.g. slight tilt of imaging field) and biological properties (e.g. vasculature) may produce inhomogeneities in rate/intensity that could be mistaken for clustering. The L function estimates the expected number of discovered neurons for different diameter circular search areas centered on each neuron, given the modeled Poisson process. We corrected for windowing in the selected field-of-view using Ripley’s isotropic correction. Global envelopes were generated (using the *envelope* function) with p < 0.05, Bonferroni-corrected.

For 2-photon opsin measurements, for each field of view we corrected for neuropil signal by manually selecting a region of neuropil with no visible cell bodies/processes and subtracting that intensity. We measured red fluorescence in donut-shaped regions of interest around the border of each cell mask. Each animal’s distribution of opsin was normalized by dividing by their maximum opsin fluorescence, and then combined. We fit a lognormal distribution via least-squares (details in Fig. S6).

### Analysis of widefield imaging data

Widefield fluorescence images were motion corrected for rigid translation, and any linear trend across the full imaging session was estimated via regression and subtracted. Deconvolution was done via Widefield Deconvolution^61^, which differs from single-neuron deconvolution algorithms like OASIS by dropping the sparsity assumption useful for spike trains of single neurons. This algorithm produces better results for aggregated signals, such as that from a single pixel during widefield imaging^61^. We rescaled the deconvolved signals to the maximum dF/F of the imaging data, as with the two-photon deconvolution. Comparison of Widefield Deconvolution, OASIS, and first-differencing is given in Fig. S3. For timecourse analyses, center and surround ROIs were defined as as the top 30% of elevated or suppressed pixels within a 1 mm radius of the center of response. To average images across animals, images were aligned on the basis of their maximum response during the late stimulation period. For quantification of spatial falloff (Fig 4G–I), we found the peak, averaged the responses radially, and then fit a curve to the responses vs. distance (LOWESS; 95% CI via bootstrap). Crossing points are the minimum distance at which the 95% confidence interval contains zero.

### Spiking network model

We simulated a conductance-based neural network model with 10000 neurons (8000 excitatory, 2000 inhibitory) to understand the recurrent features that contribute to the response properties we observe during excitatory cell stimulation. Simulations were performed using Brian2^62^.

Membrane and synaptic dynamics evolve according to the following equations:

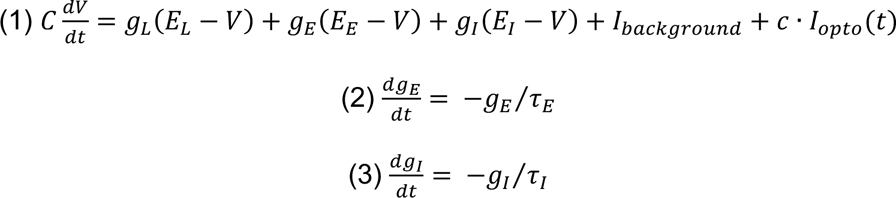

Each synapse was stepped by its corresponding connection weight for every presynaptic spike.

Connections between neurons were made with 2% probability, independently for each potential connection^36,41,63^.

Synaptic weights were drawn from truncated (rectified) Gaussian distributions. Mean connectivity parameters were based on published measurements, with excitatory connection strength an order of magnitude weaker than inhibitory connection strength^36,64,65^ and I-to-E connectivity stronger than I-to-I connectivity^65^. Background currents were chosen for inhibitory and excitatory cell populations to fix baseline firing-rates for each constructed network to data (Fig S8F). Network parameters shown in Table 1.

**Table 1:**
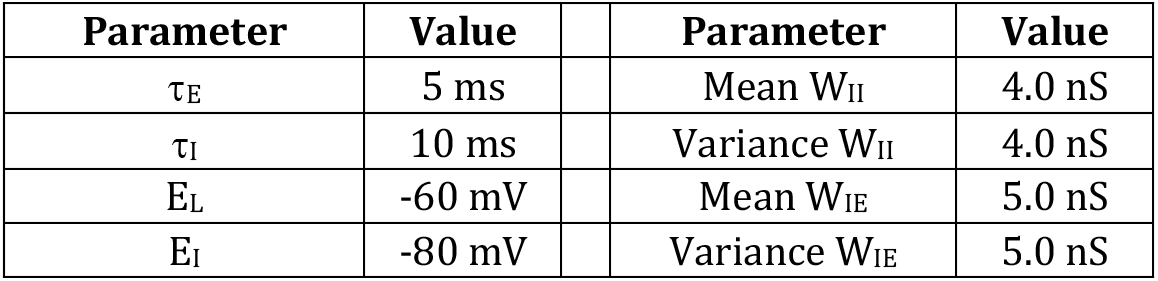
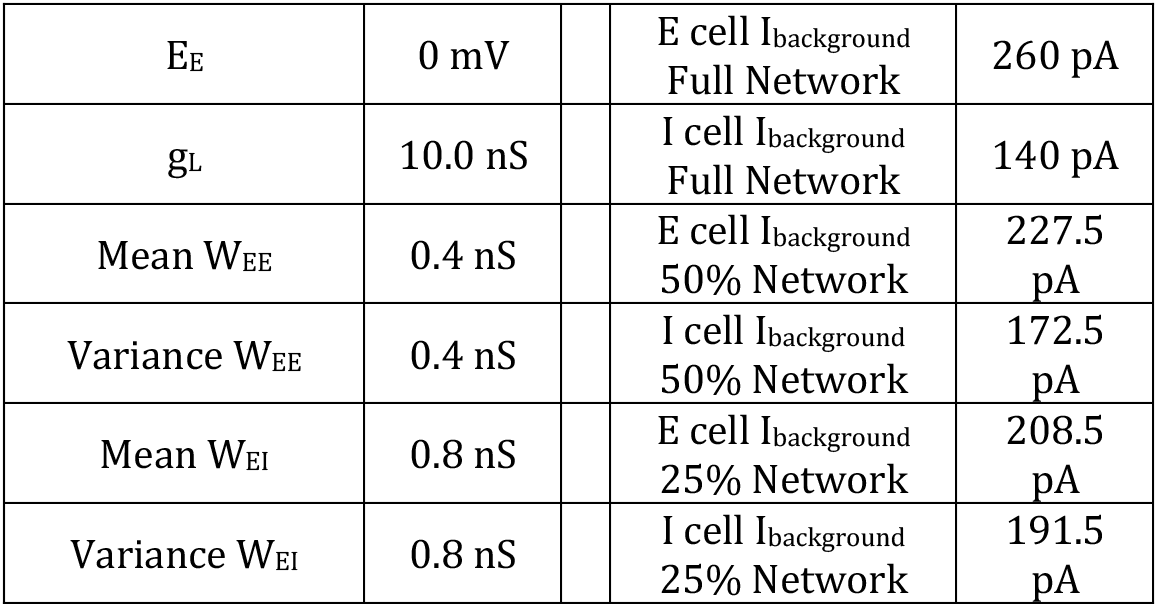
Spiking neural network model parameters.

| Parameter | Value |  | Parameter | Value |
| --- | --- | --- | --- | --- |
| $\tau_E$ | 5 ms | | Mean $W_{II}$ | 4.0 nS |
| $\tau_I$ | 10 ms | | Variance $W_{II}$ | 4.0 nS |
| $E_L$ | -60 mV | | Mean $W_{IE}$ | 5.0 nS |
| $E_I$ | -80 mV | | Variance $W_{IE}$ | 5.0 nS |

**Table 1: Spiking neural network model parameters**
|  |  |  |  |
| --- | --- | --- | --- |
| $E_E$ | 0 mV | E cell $I_{\text{background}}$<br>Full Network | 260 pA |
| $g_L$ | 10.0 nS | I cell $I_{\text{background}}$<br>Full Network | 140 pA |
| Mean $W_{EE}$ | 0.4 nS | E cell $I_{\text{background}}$<br>50% Network | 227.5 pA |
| Variance $W_{EE}$ | 0.4 nS | I cell $I_{\text{background}}$<br>50% Network | 172.5 pA |
| Mean $W_{EI}$ | 0.8 nS | E cell $I_{\text{background}}$<br>25% Network | 208.5 pA |
| Variance $W_{EI}$ | 0.8 nS | I cell $I_{\text{background}}$<br>25% Network | 191.5 pA |

### Optogenetic stimulation simulations

Optogenetic stimulation was an additional constant current for the length of the stimulation period, with onset and offset ramped linearly over 3 ms. The strength of the optogenetic stimulation (c in Eq. 1 was chosen from a lognormal distribution derived from data (Fig. 6), or held constant (Fig. 7M,N). For each simulation, this stimulation distribution was scaled by a constant to reproduce the response rate from data, at the 75^th^ percentile across excitatory cells. Steady-state response was measured for each cell as their firing rate during the 1 s baseline period subtracted from the firing rate during the last 500 ms of the stimulation period. To reduce connection strength variability (Fig. 7N), we reduced the variability of the truncated Gaussians that define connection strength by a factor of 100 (setting both the synaptic strength variability and connection number variability to zero produced a network that was less stable).

### Single cell stimulation simulations

A single cell was stimulated with intensity from maximum of input distribution (Fig. 7H). Controls: same parameters but no stimulation. To simulate single cell stimulation with visual input, we provided single cell stimulation during either the lognormal optogenetic stimulation, as previously described, or during uniform input of both excitatory and inhibitory cells.

### Balance index

We computed the balance index as described by Ahmadian and Miller (2021). For each neuron, we computed this index as the net current (excitatory + inhibitory) divided by the excitatory current. The index becomes smaller as balance becomes tighter, with component currents becoming larger, and the index becomes larger as inhibitory input from the network shrinks.

### Quantification and Statistical Analysis

All analyses, unless specifically noted in Methods Details, were performed in python using NumPy and SciPy packages^57,66^. Degrees of freedom and statistical tests are described in the results text. Error metrics plotted in figures are listed in the figure legends. Significance was adjusted for multiple testing using a Bonferroni correction when appropriate.

## Supplemental Figures

**Figure S1:**
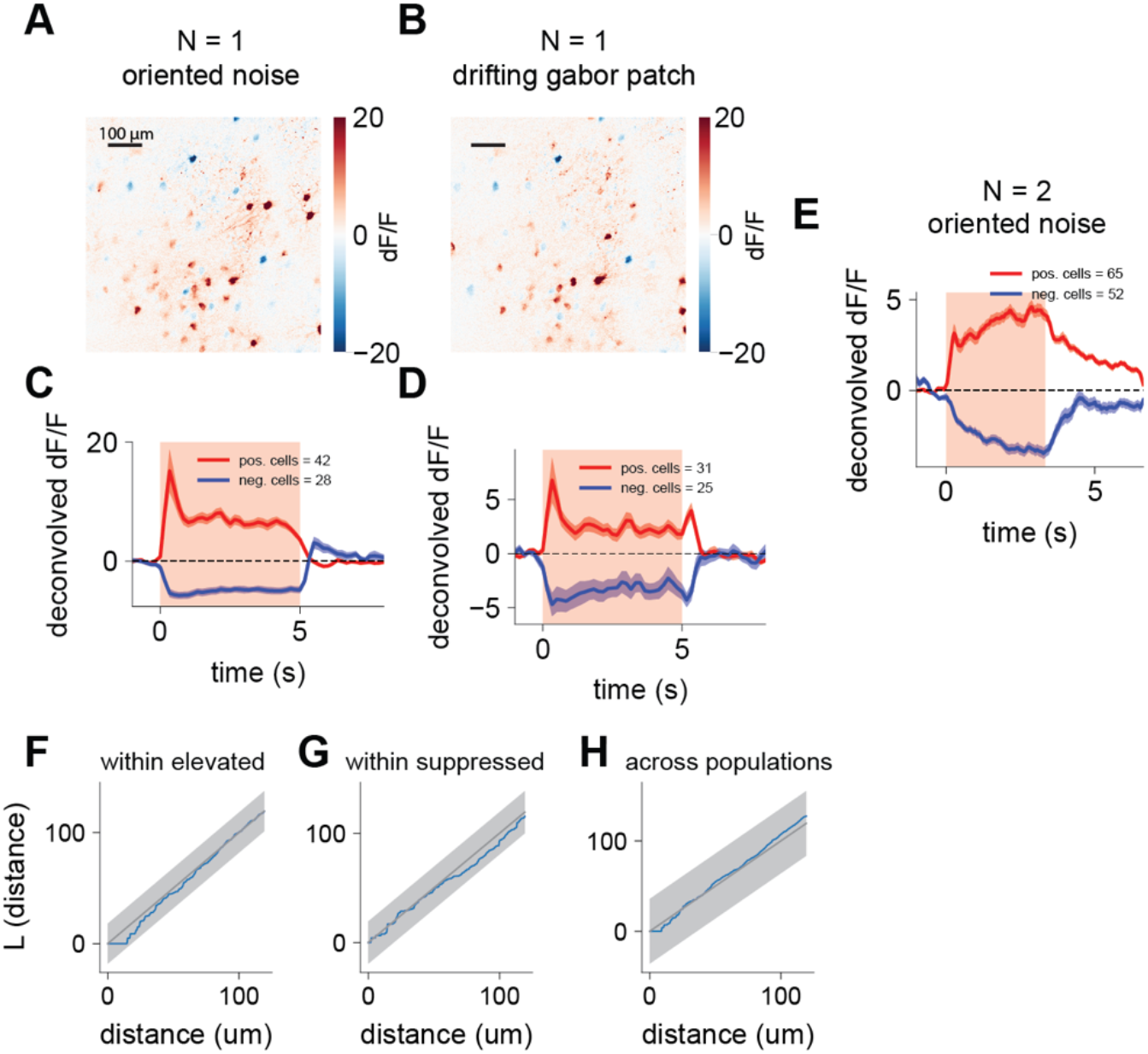
(A-E): Both grating patches and oriented noise stimuli produce steady-state elevation and suppression in layer 2/3 of V1. **(A)** dF/F response at each imaging pixel to oriented noise stimuli (small stimulus, FWHM = 15 deg, stimulus approximately aligned to cells’ receptive fields measured outside this experiment, same animal as in Fig. 1), corresponding to the deconvolved cell responses shown in Fig. 1B. Here and in Fig. 1B, responses are measured beginning 750 ms after stimulus onset to focus on steady-state response (Methods.) Evidence of suppression is seen here but is more evident when data is deconvolved (compare this panel to Fig. 1B), as expected for sustained suppression preceded by a transient, as the initial transient seen in Figs. 2, 3, 7. **(B)** dF/F response to a drifting grating (Gabor patch, spatial frequency 0.1 cpd, FWHM 15 deg), from the same animal, showing cells that are elevated and suppressed in response to drifting gratings. Overall pattern of responses to noise stimulus and grating is similar. **(C)** Deconvolved population response to oriented noise stimulus (replicated from Fig. 1F for comparison.) Stimulus on during time indicated by light red shaded box. **(D)** Deconvolved cell responses to Gabor patch, same data as in (B). Gabor patches drive both steady-state elevation and suppression, though show signs of stronger off responses and potentially a larger onset transient. **(E)** Population deconvolved response to oriented noise stimulus in two additional animals, consistent with effects from example animal. **(F-H): Spatial distributions of elevated and suppressed cells are consistent with an inhomogeneous spatial Poisson process, independent within and across classes. (F-H)** Example L-functions (Baddeley et al., 2015) from a typical animal (blue: data, black: expectation from Poisson process model, error bars: global envelopes of Poisson process model), showing agreement with the Poisson process model within elevated, suppressed, and across populations, respectively (all p > 0.05, Bonferroni correction). See Methods for analysis details.

**Figure S2:**
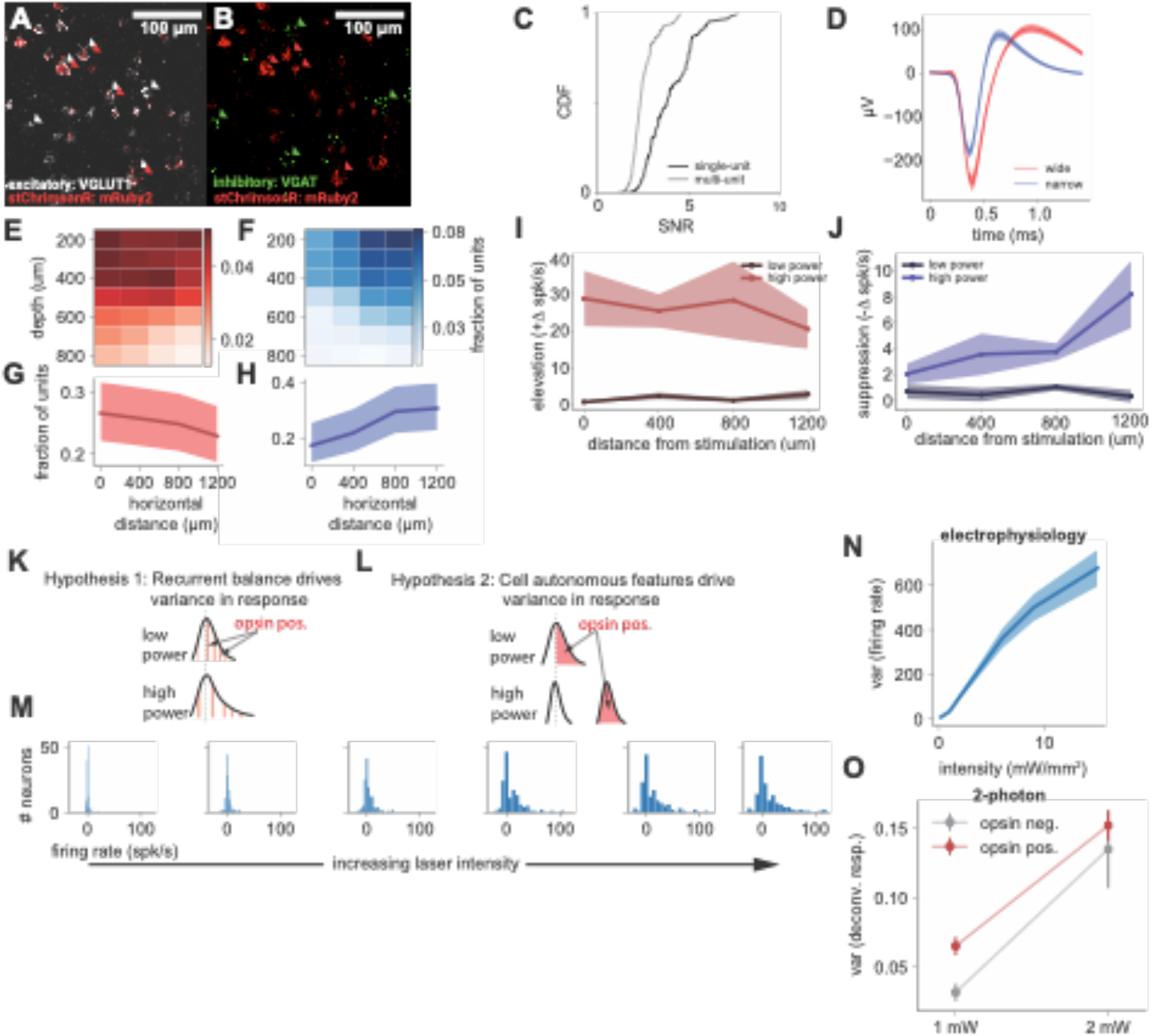
(A-B): Viral approach expresses opsin in only excitatory cells. **(A)** Selective expression of opsin in excitatory cells only, as expected for the double-inverted lox-site AAV vector and excitatory Cre mouse line (Emx1-Cre). The stChrimsonR opsin was fused to mRuby2, so we measured mRuby2 mRNA (red) and VGLUT1 mRNA (white), a marker of excitatory cells, via fluorescent in situ hybridization (RNAscope; Methods). Cell counting showed 76% of neurons are VGLUT1 positive (N = 195/257). Arrows highlight a few example neurons. As expected, all cells that express the opsin are excitatory, but not all excitatory neurons express the opsin (59% of VGLUT1 cells are mRuby2 positive: N = 115/195). **(B)** mRuby2 mRNA (red) and VGAT mRNA (green), a marker of inhibitory neurons. 24% of neurons are VGAT positive (N = 62/257), and zero express the opsin. **(C-D): Sorting and quality of electrophysiology data (C)** Single units demonstrate higher SNR (N=136, median = 3.32) than multi-units (N=184, median = 2.26). **(D)** Mean spike-waveform of putative excitatory units (wide) in red (N = 94), mean spike-waveform of putative inhibitory units (narrow) in blue (N = 42.) Bimodal histogram of spike widths is shown in Fig. 5B. **(E-H): Number of detected elevated and suppressed units by depth and horizontal distance, presented in terms of proportion of the units in the population. (E-F)** Fraction of neurons found at each depth and horizontal distance for elevated (red) and suppressed (blue) neuron populations. **(G-H)** Same as A-B, but summed across depth. Error bars: Wilson score 95% CIs. **Steady-state firing rates of neurons in layer 2/3 follow a weak spatial gradient with similar trends as the spatial distribution observed in cell counts. (I)** Elevated cell steady-state rates, with the highest and lowest powers for comparison. Rate is the difference in firing rate during stimulation relative to baseline. **(J)** Suppressed cell steady-state rates, with the highest and lowest powers for comparison, measured in relation to decreases from baseline. **(K-O): Shape and variance of response distributions are inconsistent with cell-autonomous effects. (K)** Competing hypotheses for response distribution shape. If the variance in responses is driven by network input, we would expect that responses would not be strongly correlated to opsin expression levels, and also as stimulation increases, response variance would also increase. **(L)** If cell-autonomous features like opsin expression levels drive the responses at high powers, the opsin input should dominate network input, leading to variance decreases and/or a bimodal response distribution. **(M)** Histograms of the electrophysiological response for increasing laser intensities. **(N)** Variance of the distributions in (A), plotted across laser intensity. Shaded blue: standard error. **(O)** Two-photon response variance to optogenetic stimulation, sorted by estimates of opsin expression. We see an increase in variance in both the opsin positive and negative cells, which does not support the cell-autonomous account.

**Figure S3:**
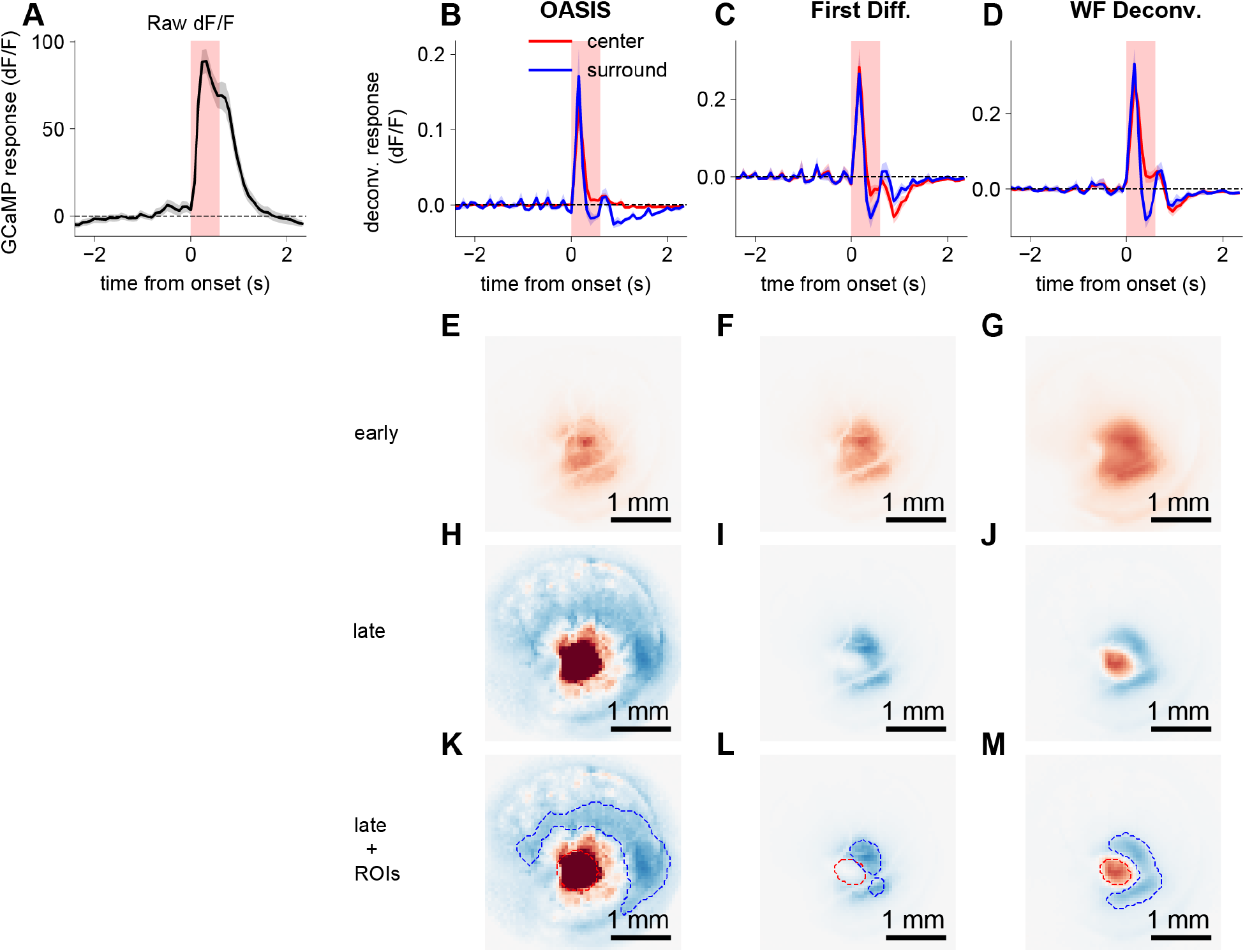
Center-surround organization is present regardless of deconvolution method. **(A)** Mean whole-frame dF/F GCaMP response in an example animal. **(B)** We tested 3 different methods of deconvolution, OASIS ^59^, first-differences (i.e. subtracting one frame from the previous), and Widefield Deconvolution ^61^. Widefield Deconvolution is expected to be the best method, as it is designed for data like this and does not incorporate the sparse-event constraints of OASIS, which is designed for single neurons. We found similar time-series results for each of the methods. The first-differences method (i.e. deconvolution with an kernel that decays immediately) seems to overestimate decreases in firing rate, as might be expected. All panels use the same dataset. **(E, F, G)** Spatial distribution of responses during the early laser period. All deconvolution methods produce a qualitatively similar excitatory response during this early period. **(H, I, J)** Spatial distributions of responses during late laser period demonstrates slight differences in size of surround, but overall a qualitatively similar center-surround organization with all methods. **(K, L, M)** Spatial distributions of response during the late laser period, but with dashed contours depicting the manually-drawn regions of interest (ROIs) that we used to produce the time-series data in **(B, C, D)**, with red dashed contours representing the center ROIs, and blue dashed contours representing the surround ROIs.

**Figure S4:**
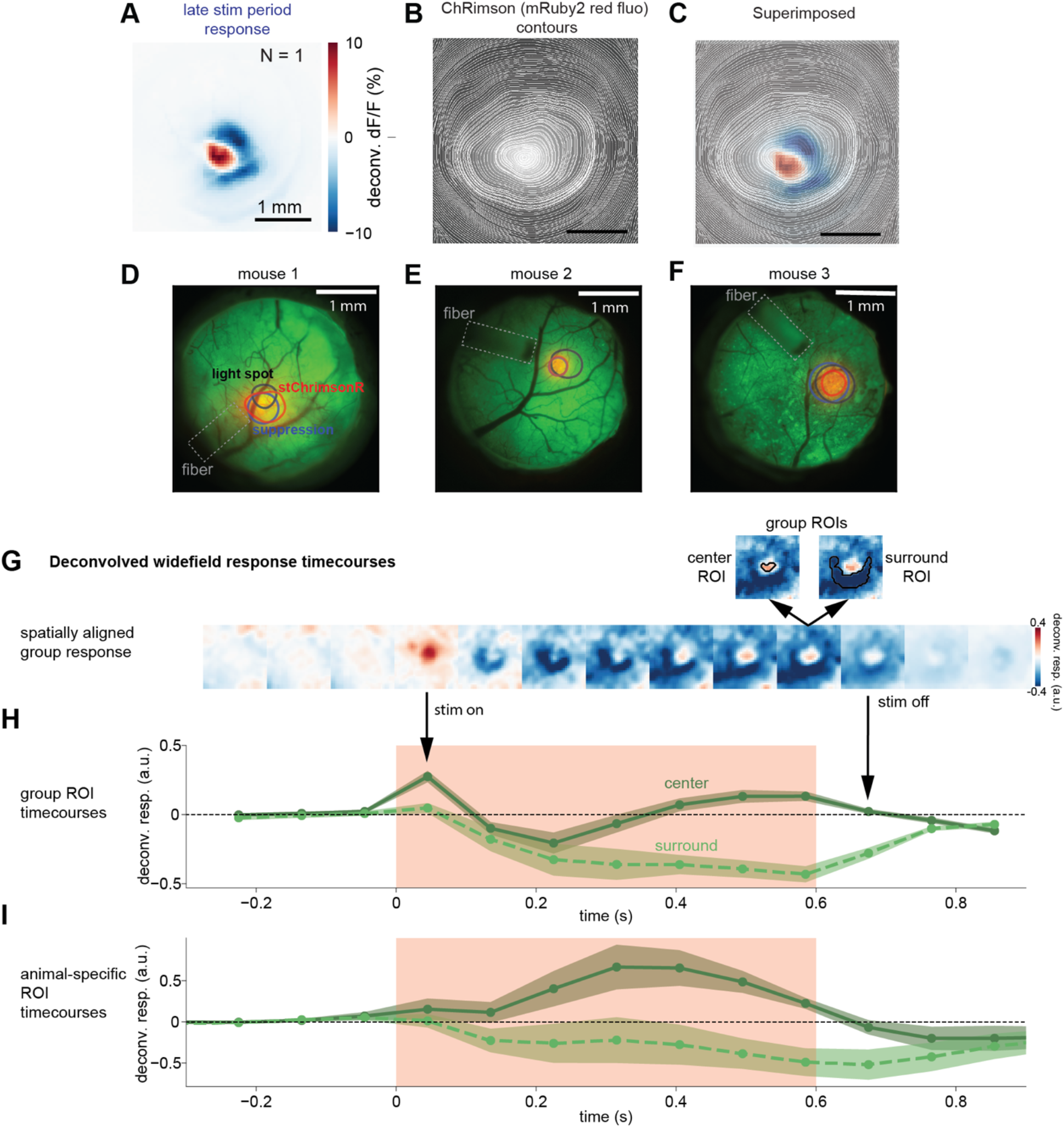
(A-C): Stimulation response correlates to the pattern of stChrimsonR expression. **(A)** Example animal’s response during the late stim period. **(B)** Example animal’s stChrimsonR expression pattern (gray: fluorescence) with overlaid contours of fluorescence intensity. **(C)** Example animal’s response during the late stim period overlaid with their stChrimsonR contours. **(D-F): Expression and surround response of each mouse in the widefield dataset. (D-F)** Field-of-view showing GCaMP expression (green image), stChrimsonR expression (red image). Contours: black = 80% of maximum illumination, red = 80% of maximum expression, blue = local minimum of the surround suppression. **(G-I): Spatiotemporal response pattern of widefield response to excitatory cell stimulation. (G)** Response over time, each frame corresponding to a timepoint in the timecourse in (B). Group ROIs were selected as the top 30% of positively or negatively responding pixels within 1 mm of the center of response and were used to compute the timecourses in (B); Methods. **(H)** Timecourse of the response in the center and surround in the group-averaged signal. Error: standard deviation across pixels. **(I)** Reproduction of Fig. 4D. Same as (B), but each animal’s timecourse was generated from their individual data and then averaged, resulting in less smoothing between center and surround due to small variations in optogenetic expression region size across animals. Error: standard error across animals.

**Figure S5:**
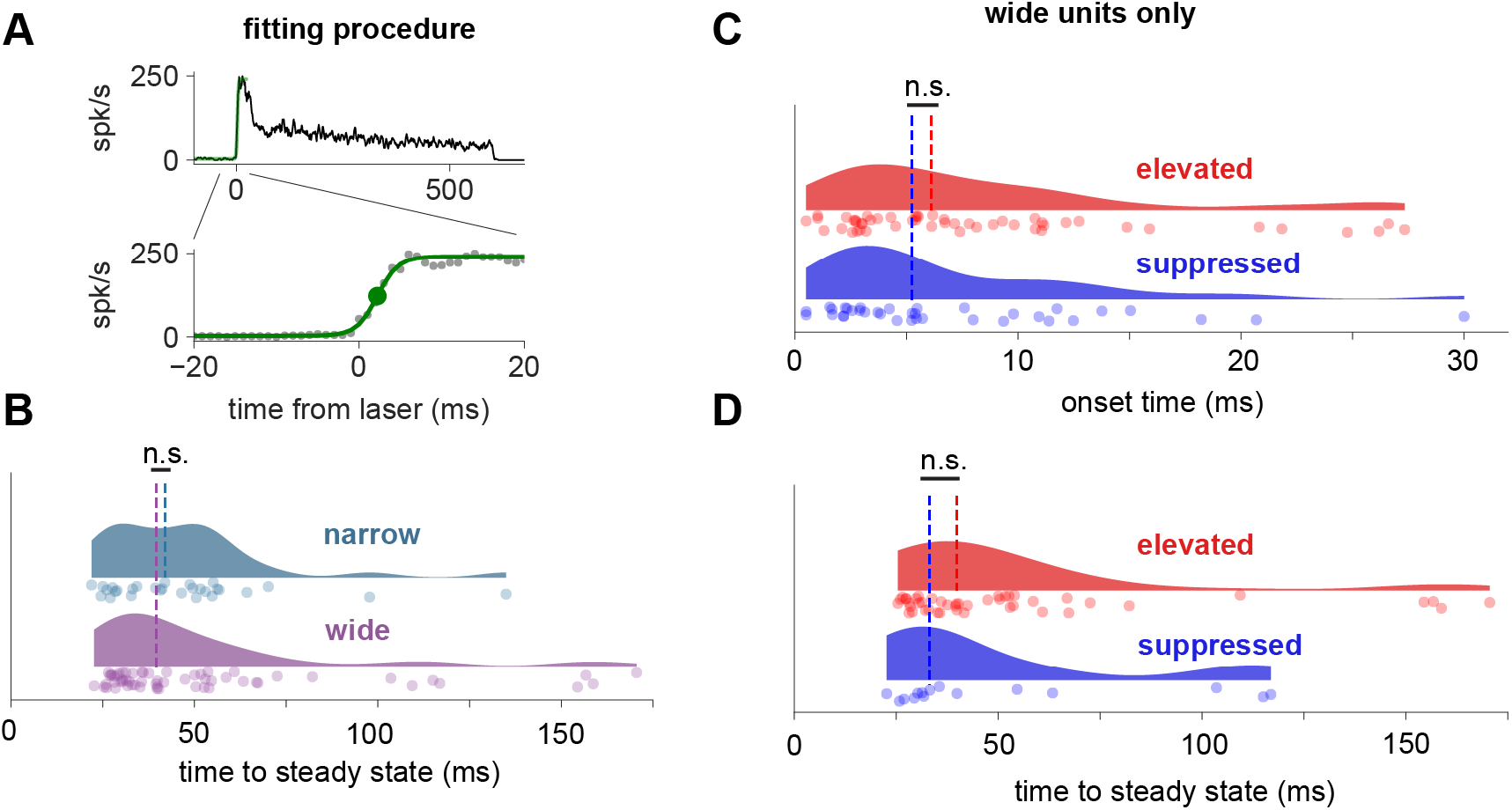
Differences in dynamics are restricted to those seen between the onset of wide- and narrow-waveform cells. The excitatory and inhibitory (wide- and narrow-) onset latency difference is shown in Fig. 7C. Other quantities shown here do not differ: wide-vs narrow (excitatory vs inhibitory) time to steady state (B), and onset time and time to steady state (C,D) for elevated and suppressed groups of wide-waveform excitatory cells. **(A)** Example single neuron firing rate with fits. To obtain the onsets for individual cells, each cell’s mean timecourse was smoothed with width dependent on the detectability of the transient signal (SNR; Methods), then a logistic function was fit to data from time range [-100ms, 100ms]. The onset time (latency) was defined as the time to half-max of the logistic function. **(B)** No difference in median time to steady state was found across narrow-spiking and wide-spiking cells. **(C)** No difference in median onset time for elevated and suppressed groups of wide-waveform (excitatory) cells. **(D)** Same as B, but difference in median time to steady-state.

**Figure S6:**
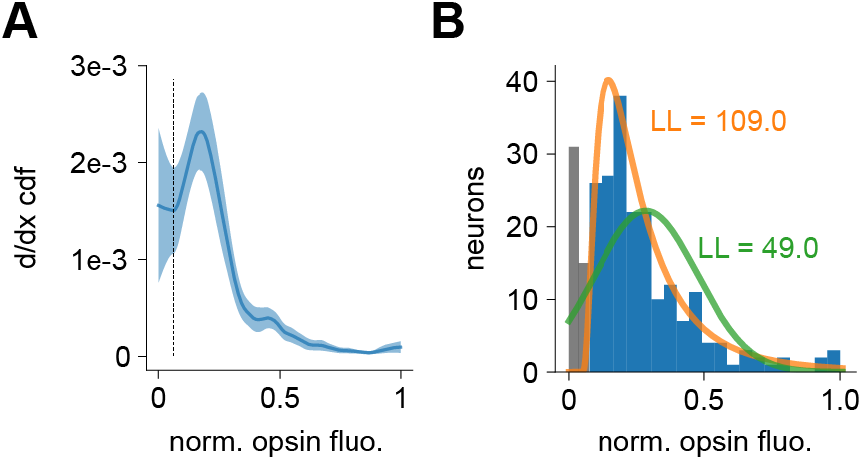
Distributions of opsin fluorescence measured in vivo. **(A)** Red channel after neuropil correction (Methods). Y-axis, first derivative of CDF, smoothed with LOWESS; point separating non-expressing neurons (left, below dashed line) and expressing (above dashed line) is set at the local minimum. Error bars (light blue): bootstrapped standard error (N=244 neurons, N=3 animals). **(B)** Histogram, same data. The log-likelihoods (LLs) indicate that a lognormal distribution (orange) fits the observed distribution better than a Gaussian (green). Shown: fits used for simulations, excluding non-expressing neurons (gray). LLs, lognormal = 109.0, Gaussian 49.0. (LLs when including all neurons: lognormal = 95.3, Gaussian = 47.5).

**Figure S7:**
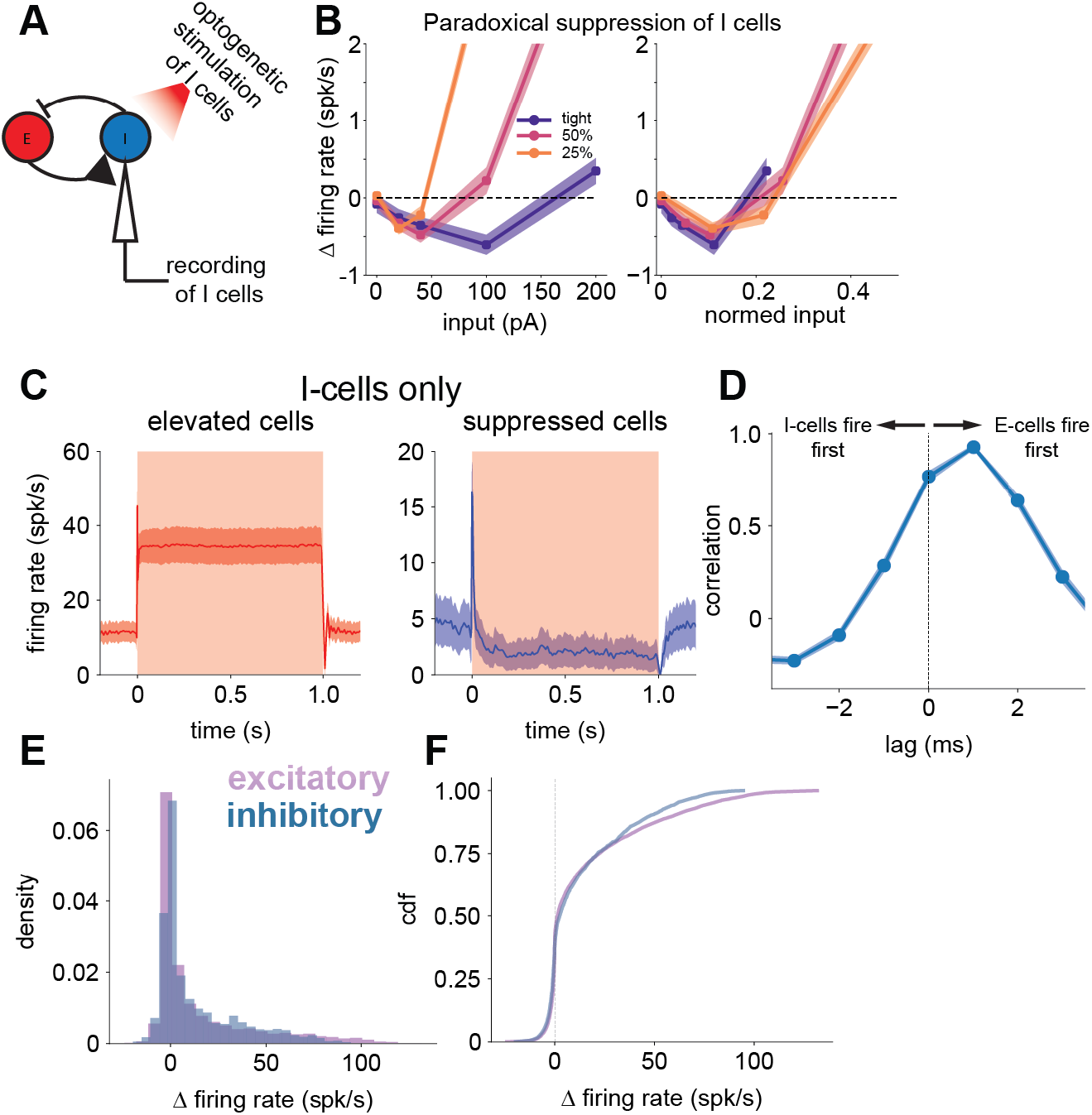
(A-B): Networks at all tested recurrent strengths operate within the ISN regime. **(A)** To examine paradoxical suppression, we record the steady-state responses of the I cells in response to different levels of stimulation. We performed this experiment on all networks presented in Figure 8. **(B left)** Steady-state responses of the I cell population to 5 levels of stimulation. Error bars are standard error to the mean. Graph has been zoomed into the region which clearly shows paradoxical suppression in all 3 networks. This paradoxical suppression is predicted for both loosely and tightly balanced networks. Our simulations used three recurrent strength values, one in the tight-balance regime and two in the loose-balance regime, and we confirmed that all three showed paradoxical effects of suppression when I cells are stimulated **(B right)** Same as (B left) but input normalized by the input value calculated in Fig. 8 to drive each network to the same firing rate (input level that achieves same value of the 75^th^ percentile of evoked rates; see Fig. 8). **(C-F): Inhibitory neurons in balanced state model show similar responses to excitatory neurons but are recruited after initial stimulation. (C)** Mean timecourses for elevated and suppressed inhibitory cells (left and right, respectively) show the same characteristic transient response followed by steady-state responses. **(D)** Cross-correlation analysis of E- and I-cell response. Network has no synaptic delays built into the model. E-cells respond to direct stimulation, and then I-cells are recruited after. **(E)** Population distribution of steady-state responses is similar across E- and I-cells, though excitatory cells show a slightly longer-tailed positive response (true in the data as well; Fig. 5E), as seen through the distribution of responses or their corresponding CDFs **(F)**

**Figure S8:**
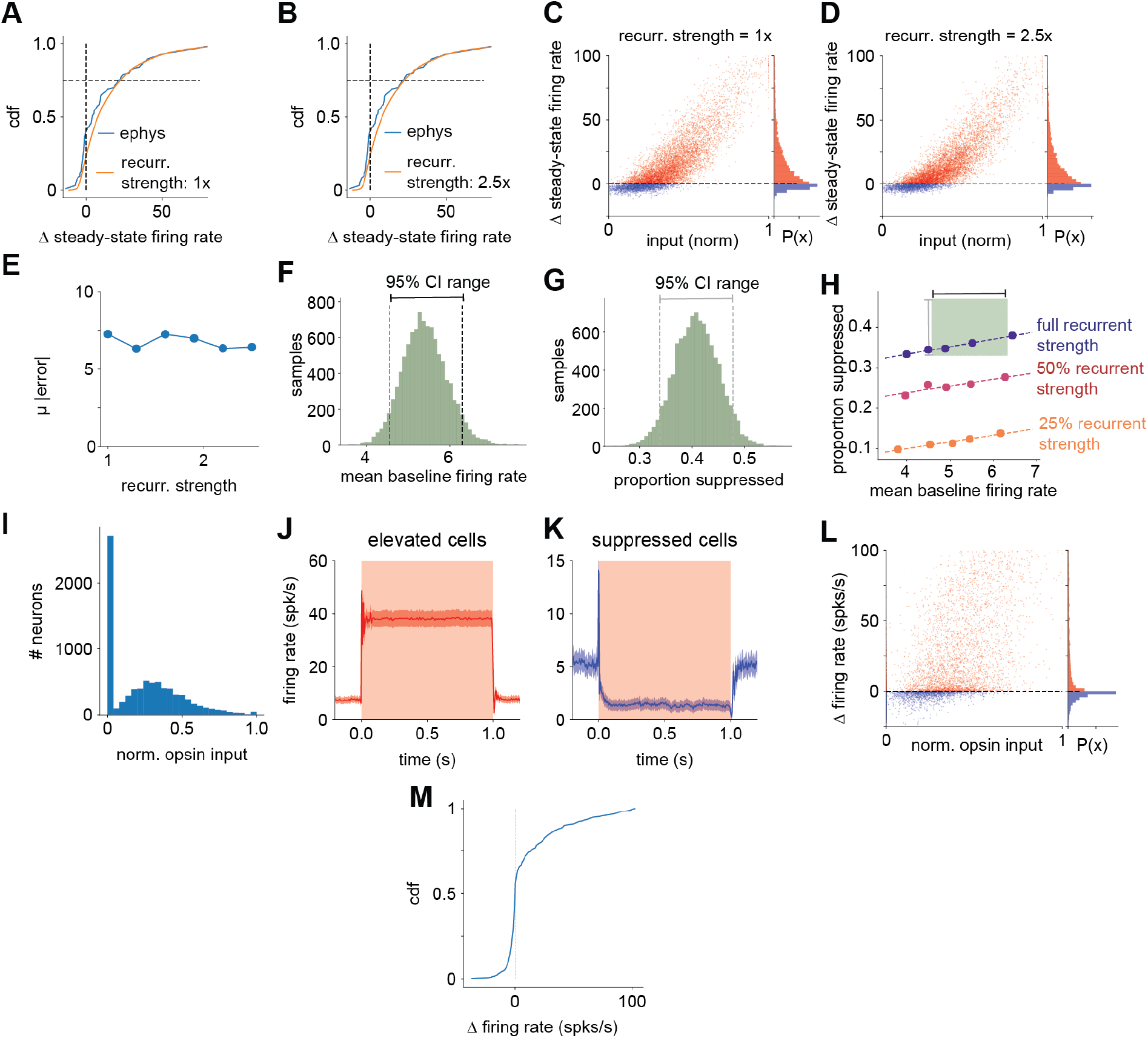
(A-E): Increasing strength of recurrent connections does not substitute for recurrent connection variability. **(A)** Cumulative distribution of responses to optogenetic stimulation in model with 1x recurrent strength, matched to the 75^th^ percentile of the response measured using electrophysiology. Negligible recurrent variability in this simulation (same number of recurrent connections to each neuron, variability in recurrent strength ∼1% of mean, see Methods), and so spread in responses as a function of input is due to optogenetic input variability. Distribution of input is inferred from data in Fig. 6 (lognormal fit; Methods.) **(B)** Same as (A) but in model with 2.5x recurrent strength. **(C)** Relationship between input and steady-state response in the model with 1x recurrent strength. Marginal distribution of response show on the right. **(D)** Same as (C), but in model with 2.5x recurrent strength. Note that both stimulations produce similar variability between input strength and firing rates. This variability is seen as spread in the red cloud of points around an imagined curve that could be fit through the points. **(E)** Estimated mean absolute error of the relationship between the input and output as measured by a LOWESS fit across all recurrent strength manipulations. **(F-H): Sensitivity analysis demonstrates that matching suppression is achievable within confidence bounds observed baseline firing rates, but only in the network with the strongest recurrent connectivity strength. (F)** Bootstrapped distribution of baseline firing rate estimated from electrophysiology data. 95% confidence intervals are drawn from the bootstrapped distribution. **(G)** Bootstrapped distribution of the proportion of the population that is suppressed following optogenetic stimulation, estimated from the electrophysiology data. **(H)** The network baseline firing rate and recurrent strength were systematically manipulated, finding that the only networks that can replicate the proportion of suppression we observe within the baseline firing rate we observe are networks with strong recurrent connectivity. **(I-M): Models with 41% of cells without opsin replicate steady-state dynamics, noisy relationship between opsin input and steady-state response, and response distribution. (I)** Distribution of opsin input was generated by sampling from a lognormal distribution fit to our observed opsin fluorescence, and in order to replicate the sparse expression we observed in histology we set 41% of cells to 0 at random. **(J)** Mean timecourse of response to stimulation in elevated cells maintains the same transient and steady state dynamics observed in the main simulations. **(K)** Same as (B), but in suppressed cells. **(L)** Relationship between opsin input and steady state response remains weak but positive, marginal distribution shown on right. **(M)** Cumulative response distribution to stimulation shows typical long tail and large proportion of suppressed responses.

